# Analytical guidelines to increase the value of citizen science data: using eBird data to estimate species occurrence

**DOI:** 10.1101/574392

**Authors:** A Johnston, WM Hochachka, ME Strimas-Mackey, V Ruiz Gutierrez, OJ Robinson, ET Miller, T Auer, ST Kelling, D Fink

**Affiliations:** Cornell Lab of Ornithology, Cornell University, 159 Sapsucker Woods Road, Ithaca, NY, 14850, USA

**Keywords:** citizen science, detectability, eBird, encounter rate, occupancy model, spatial bias, species distribution model

## Abstract

Citizen science data are valuable for addressing a wide range of ecological research questions, and there has been a rapid increase in the scope and volume of data available. However, data from large-scale citizen science projects typically present a number of challenges that can inhibit robust ecological inferences. These challenges include: species bias, spatial bias, and variation in effort.

To demonstrate addressing key challenges in analysing citizen science data, we use the example of estimating species distributions with data from eBird, a large semi-structured citizen science project. We estimate two widely applied metrics of species distributions: encounter rate and occupancy probability. For each metric, we assess the impact of data processing steps that either degrade or refine the data used in the analyses. We also test whether differences in model performance are maintained at different sample sizes.

Model performance improved when data processing and analytical methods addressed the challenges arising from citizen science data. The largest gains in model performance were achieved with: 1) the use of complete checklists (where observers report all the species they detect and identify); and 2) the use of covariates describing variation in effort and detectability for each checklist. Occupancy models were more robust to a lack of complete checklists and effort variables. Improvements in model performance with data refinement were more evident with larger sample sizes.

Here, we describe processes to refine semi-structured citizen science data to estimate species distributions. We demonstrate the value of complete checklists, which can inform the design and adaptation of citizen science projects. We also demonstrate the value of information on effort. The methods we have outlined are also likely to improve other forms of inference, and will enable researchers to conduct robust analyses and harness the vast ecological knowledge that exists within citizen science data.

## Introduction

Citizen science data are increasingly making important contributions to basic and applied ecological research. One of the most common forms of citizen science data comes from members of the public recording species observations. These observations are being collected for a diverse array of taxa, including butterflies (Howard, Aschen, & Davis, 2010), sharks (Vianna, Meekan, Bornovski, & Meeuwig, 2014), lichen (Casanovas, Lynch, & Fagan, 2014), bats (Newson, Evans, & Gillings, 2015), and birds (Sauer et al., 2017). The number of these citizen science projects has been growing exponentially, but they vary widely in complexity, flexibility, and participation (Pocock, Tweddle, Savage, Robinson, & Roy, 2017; Wiggins & Crowston, 2011). Projects occur on a spectrum from those with a predefined sampling structure that resemble more traditional survey designs, to those that are unstructured and collect observations ‘opportunistically’. Projects with study designs and defined protocols generally produce data that are more informative for a particular objective, but are often limited to a specific time frame and region and have fewer participants. This can lead to a trade-off between the quality and quantity of data supported by citizen science projects (Bird et al., 2014; Pacifici et al., 2017). Semi-structured citizen science projects have unstructured data collected, but critically also collect data on the observation process, which can be used to address many of the issues arising with citizen science data (Altwegg & Nichols, 2019; Kelling et al., 2018). With the increasing popularity in the use and application of citizen-science data, we describe and evaluate steps for data processing and analysis that maximise the value of semi-structured citizen science data (Sullivan et al., 2014).

Data consisting of species observations from citizen scientists present a number of challenges that are not as prevalent in conventional scientific data. Firstly, participants often have preferences for certain species, which may lead to preferential recording of some species over others (Troudet, Grandcolas, Blin, Vignes-Lebbe, & Legendre, 2017; Tulloch & Szabo, 2012). Secondly, the observation process is heterogeneous, with large variation in effort, time of day, observers, and weather, all of which can affect the detectability of species (Ellis & Taylor, 2018; Oliveira, Olmos, dos Santos-Filho, & Bernardo, 2018). Thirdly, the locations selected by participants to collect data usually are strongly spatially biased. For example, participants may preferentially visit locations that are close to where they live (Dennis & Thomas, 2000; Mair & Ruete, 2016), more accessible (Botts, Erasmus, & Alexander, 2011; Kadmon, Farber, & Danin, 2004), contain high species diversity (Hijmans et al., 2000; Tulloch, Possingham, Joseph, Szabo, & Martin, 2013), or are within protected areas (Tulloch et al., 2013). However, citizen science data also contain a wealth of ecological information and they are often the only source of biological knowledge for many biodiverse regions. Therefore it is imperative to define approaches that can maximise the value of increasing volumes of citizen science observations.

There are two main approaches for addressing known challenges related to citizen-science data: 1) imposing a more structured protocol onto the dataset after collection via data filtering (Kamp, Oppel, Heldbjerg, Nyegaard, & Donald, 2016); 2) including covariates in a model to account for the variation (Miller, Pacifici, Sanderlin, & Reich, 2019). In this paper we advocate combining both of these approaches to increase the reliability of inferences made using citizen science observations.

We describe analytical approaches for using semi-structured citizen science data, using the example of estimating species distributions from data collected by the eBird citizen science project (Sullivan et al., 2014). We use two critical aspects to these citizen science data that facilitate robust ecological inference. Firstly, data submitted to eBird are structured as ‘checklists’, where each checklist is a list of bird species recorded during one period of bird-watching. Secondly, eBird is a semi-structured citizen science project, which means most eBird checklists have associated metadata describing the ‘effort’ or observation process (Kelling et al., 2018). While our examples focus on the use of eBird data, our results are important both for analysis of similar citizen science datasets, and for the design of future citizen science surveys.

## Methods

We explored the impact of various analytical practices when using citizen science data to estimate species distributions. We used different modelling approaches to estimate: 1) encounter rate with Maxent and a random forest model, and 2) occupancy rate with an occupancy model. Species encounters arise as a compound process requiring both the species to occur at a site and to be detected at that site. Encounter rate is defined as the average rate at which observers encounter the species, so it reflects the product of occurrence and detectability. Occupancy is defined as the probability that a species is present in a given location, because the model separates occurrence and detectability. While Maxent uses only detection (or “presence-only”) data, random forest and occupancy models use detection/non-detection data as their response variable. All analyses were conducted with R (R Core Team, 2018). As a case study we focussed on wood thrush *Hylocichla mustelina* in the breeding season. Wood thrush is a relatively common passerine in north east America, that is easily detected by its song. All data and code for this analysis are within Supporting information A4 and more examples of running species distributions with eBird data are in Strimas-Mackey et al. (2020).

### eBird data selection

We used data from the eBird Basic Data (EBD), which is global in extent and updated monthly (www.ebird.org/science/download-ebird-data-products). The most current version of the EBD can be freely accessed via an online data portal and processed with the auk R package (Strimas-Mackey, Miller, & Hochachka, 2018). eBird has a robust review process, focussed on ensuring correct locations and species identification, that is conducted before data enter the EBD; we provide further details on this and other aspects of eBird data in Supporting Information A1. Our data are from the EBD version released in May 2019. To model wood thrush distribution in the breeding season, we used checklists from 15 May - 30 June. We restricted these data geographically to Bird Conservation Region 27 “Southeastern coastal plain” (BCR27), a biogeographically uniform region that includes parts of the states: Mississippi, Alabama, Florida, Georgia, North Carolina, and South Carolina (NABCI 2000).

### eBird data processing

We split the eBird data into a dataset to use to train (or fit) the models, and semi-independent datasets to validate (or test) the models (Figure 1 and S1). To train the models we used eBird data from 2018. We created two validation datasets, chosen because of their different forms of independence from these training data. The form of validation data should be tailored to a specific intent (Valavi, Elith, Lahoz-Monfort, & Guillera-Arroita, 2018). Our main validation set was temporally independent, using the using eBird data from 2017 to create a balanced validation set (equal detections and non-detections) with reduced spatial bias (Figure 1). The reduced spatial bias ensured that the data represent the study region more evenly, and the balance of detections and non-detections tested the ability of the model to discriminate between areas of species presence and absence. Our second validation dataset was the 2018 data from the North American Breeding Bird Survey that were also submitted to eBird. This created a validation against a standardized and pre-designed survey (Figure 1). See supporting information A2 for more details.

**Figure 1.**
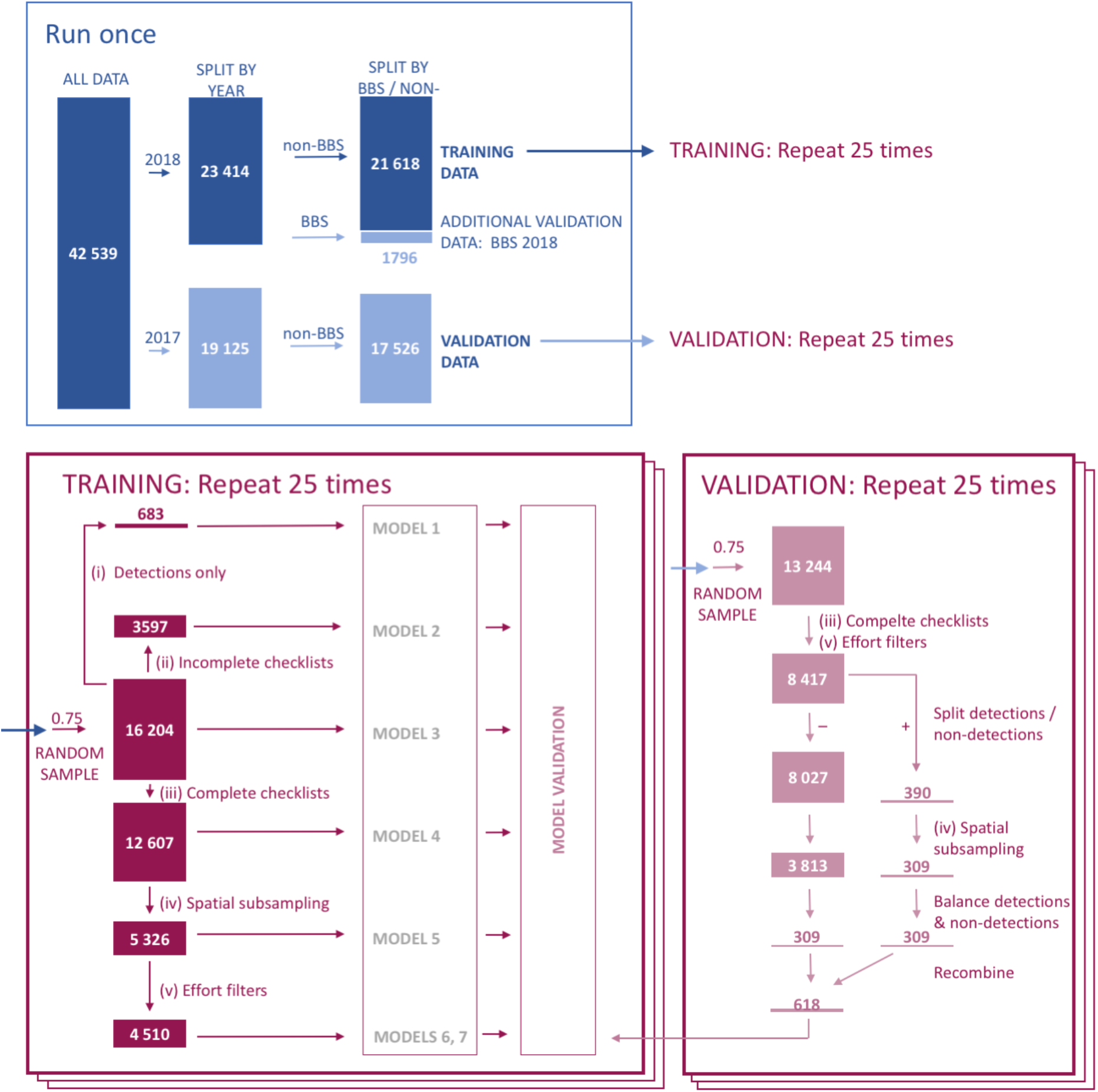
Schematic diagram of the flow of data into each of the 6 model types for the encounter rate model. The sizes of the boxes and the numbers inside them are the number of checklists. The blue processes occur once, the pink processes occur 25 times, once for each model run. The numbers shown will therefore vary slightly each time within the pink box. The dark colours represent training data and the pale colours validation data. Arrows represent data processing steps, or projection of the same data forward to the next stage.

We used a hierarchy of data processing steps on the training data, applying these sequentially to create a set of differently processed datasets. These data processing steps were designed to highlight or address the challenges with citizen science data outlined in the introduction. We applied these datasets to each of the model types to estimate both encounter rate and occupancy (Table 1). Two data processing steps were designed to degrade the data: i) select only detections; ii) select only ‘incomplete’ checklists. The models with these data (models 1 and 2) highlight the impact of having these degraded data. Three data processing steps were designed to refine the data: iii) selecting only ‘complete’ checklists; iv) spatially subsampling; and v) selecting checklists with standard effort (Table 1). Using non-detections allows the model to have knowledge of where effort was expended, but the species was not recorded. Data processing step (iii) ensures that all the inferred ‘non-detections’ are actually nondetections, by only including checklists where observers report all the species they could detect and identify (complete checklists). This addresses the challenge of species bias. Step (iv) spatial subsampling reduces the over-influence of well-surveyed locations in the analysis. This addresses the challenge of spatial bias. Step (v) reduces the range of checklist effort, creating a more consistent and standardised set of checklists for analysis. This addresses the challenge of variable effort. Methodological details for how we performed each of these data processing steps is giving in supporting information A2.

**Table 1.** Descriptions of the elements in models 1-7 that include different data processing treatments. Models 3 uses all the data with no processing. Models 1-2 use data degraded in different ways by processes (i) and (ii). Models 4-6 use data refined in different ways by processes (iii), (iv), and (v). Model 7 uses the same data as Model 6, but additionally includes effort variables as covariates.

|  | Data processing treatment |  | Model |  |  |  |  |  |  |
| --- | --- | --- | --- | --- | --- | --- | --- | --- | --- |
|  |  |  | 1 | 2 | 3 | 4 | 5 | 6 | 7 |
| DEGRADE | i) | Select detections only ('presence only') | ✓ |  |  |  |  |  |  |
|  | ii) | Select incomplete checklists only |  | ✓ |  |  |  |  |  |
| REFINE | iii) | Select complete checklists only |  |  |  | ✓ | ✓ | ✓ | ✓ |
|  | iv) | Spatial subsampling |  |  |  |  | ✓ | ✓ | ✓ |
|  | v) | Effort filters |  |  |  |  |  | ✓ | ✓ |
|  | vi) | Effort covariates |  |  |  |  |  |  | ✓ |
| MODEL TYPE | Encounter rate model type |  | Maxent | Random forest | Random forest | Random forest | Random forest | Random forest | Random forest |
|  | Occupancy model |  |  | ✓ | ✓ | ✓ | ✓ | ✓ | ✓ |

Preparing data for the occupancy models required some additional data processing. There are many decisions required when using citizen science data for occupancy models and we describe these in greater detail in the supporting information A3 with only a brief overview here. We defined a ‘site’ as a location (defined by latitude and longitude) with at least two visits during May 15 - June 30 2018. Where there were more than 10 visits to a single site, we randomly selected 10 of the visits. For the occupancy models we also created a separate validation set. We wanted to validate the estimates of occupancy, whilst limiting the effects of detectability. We therefore selected sites with high detectability and determined whether wood thrush was recorded on any visit. Using this validation dataset we compared the estimated occupancy to the observed occurrence.

### Environmental data

As environmental covariates we used land cover data derived from MODIS product MCD12Q1 v006 (Friedl & Sulla-Menashe, 2015). We estimated the land cover associated with each checklist as the proportion of each land cover category in a 2.5 km x 2.5 km square surrounding the checklist location in the year the observations were made. We included the proportions of each land cover type in the MCD12Q1 v006 classification. See Supporting Information A2 for a list of the landcover types.

### Estimating species encounter rate

We estimated the *encounter rate* of wood thrush on eBird checklists in relation to the environmental covariates. The response was the detection/non-detection of wood thrush and the environmental covariates were 16 landcover covariates as described above. Model 1 used presence-only records of wood thrush on a checklist, fitted with a Maxent model through the R package maxnet (Phillips, 2016). Models 2-7 fitted a random forest with detection/non-detection records of wood thrush on checklists, followed by calibration with a Generalized Additive Model (GAM). The random forest models were fitted with the R package ranger (Wright & Ziegler, 2017) and the GAMs within R package scam (Pya, 2013). We ran the set of seven models 25 times, and for each set we randomly selected 0.75 of the training and validation datasets before applying the relevant data processing (Figure 1). For further details of the model fitting see the supporting information A2 and the code in supporting information A4.

We validated the fitted model and calibration model with the semi-independent validation datasets. We used a range of performance metrics to compare the estimates to the observations: sensitivity, specificity, True Skill Statistic (TSS), Area Under the Curve (AUC), Kappa, and Mean Squared Error (MSE, also named Brier score). To quantify the additional benefit of running the different models, for each set of seven models, we calculated the differences in performance metrics between each model and model 3. We examined these differences across the 25 different runs of the model set.

For a random set of the seven models, we mapped the estimated encounter rate across the whole region of the BCR27. We produced a dataset with the land cover for each 2.5 km x 2.5 km grid cell across the entire region and we set effort variables to be constant across the region. The predictions relate to the hypothetical encounter rate of an average eBird participant conducting a 1 hour, 1 km complete checklist on 15 June 2018 at the optimal time of day for species detection. We used the random forest model and the calibration GAM to estimate encounter rate for this standardised checklist in each grid cell in BCR27.

### Estimating species occupancy

To assess the effects of these data processing steps in an alternative modelling framework, we applied single-species occupancy models to estimate occupancy and detectability of wood thrush. We modeled occupancy probability as a function of MODIS land cover categories (Friedl & Sulla-Menashe, 2015) and we selected four categories considered a priori to have the most ecological relevance: deciduous broadleaf forest, mixed forest, croplands, and urban. For modelling detection probability, we used the five effort covariates described in the ‘encounter rate model’ above. We used the R package unmarked to fit single-season models (Fiske & Chandler, 2011). We ran these occupancy models using a set of six different combinations of data processing and model structure (Table 1). For further details of the model fitting see the supporting information A2. We validated the occupancy predictions against the specific occupancy validation dataset of sites with high detectability. As above, we also mappedthe occupancy rate across the whole region by predicting to the whole of the BCR27.

### Varying sample size

Our study area has a relatively high density of eBird data, but other regions and other citizen science projects may often have fewer data. Therefore we wanted to assess whether the results we found would be the same smaller datasets. We estimated wood thrush encounter rate using model 3 and model 7 for a range of sample sizes. For each model set we randomly selected 0.75 of the training and validation datasets (as above). We then further reduced the data to proportions of this new total: 0.1, 0.3, 0.5, 0.7, or 0.9. We ran this set of 10 models (five sample sizes, two models) 25 times. For each run, we compared the difference in predictive performance metrics (as described above) between model 7 and model 3.

## Results

### Estimating species encounter rate

Model 7 had the highest estimates of encounter rate (Figure 2) and the best model performance. Model performance was consistently the best with model 7, across both validation datasets and most of the performance metrics (Figures 3, S2). Thus the combination of all data processing steps resulted in the best model, with adding effort variables as covariates producing the biggest improvement in our example (compare models 6 and 7).

**Figure 2.**
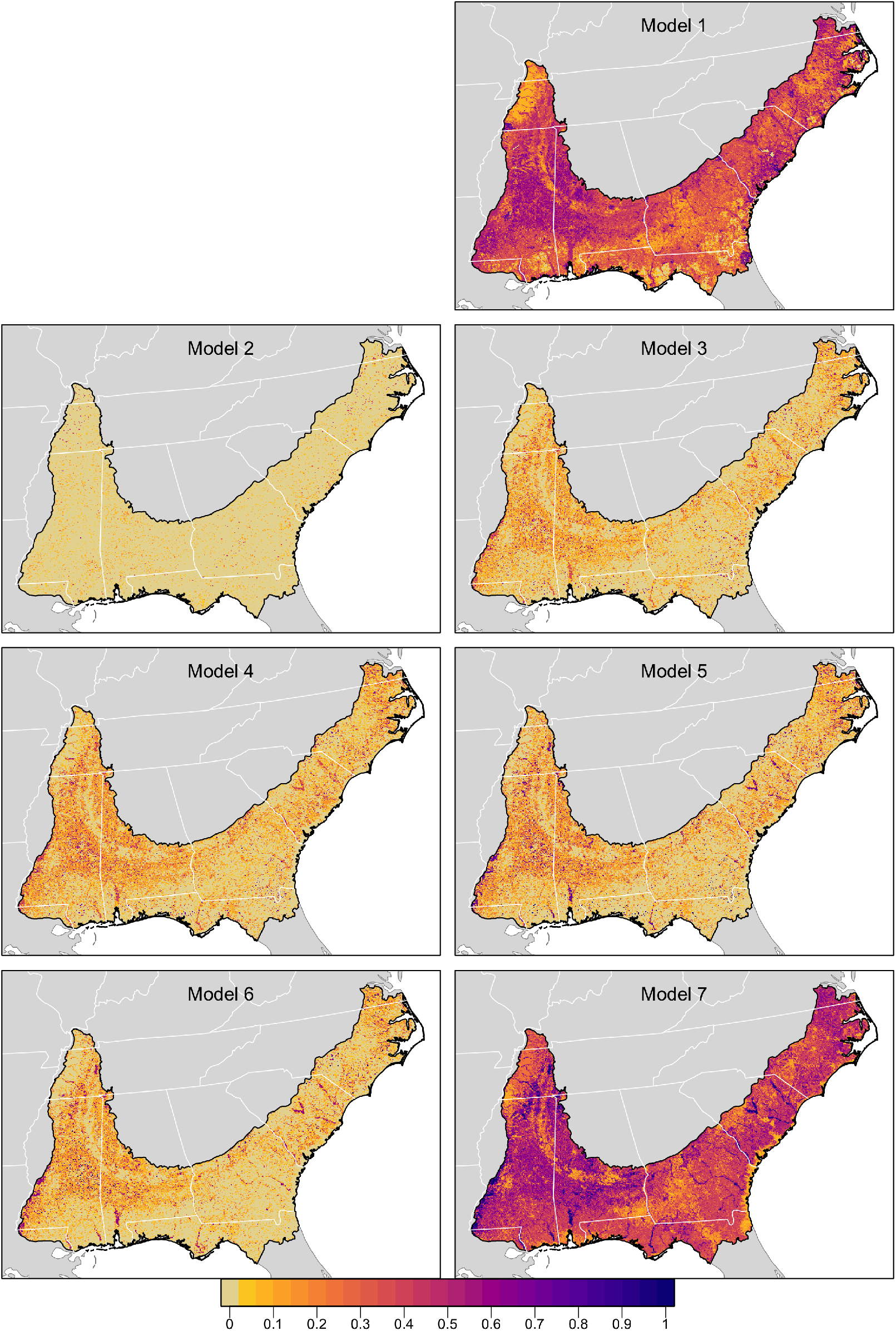
Estimated wood thrush encounter rate across the BCR27 region for models 1 - 7. Estimated encounter rate is the expected proportion of standardised checklists that would record Wood Thrush. These hypothetical standardised checklists are conducted by an average eBirder, travelling 1km over 1 hour, at the optimal time of day for detecting Wood Thrush. Darker colours denote higher estimated encounter rate.

**Figure 3.**
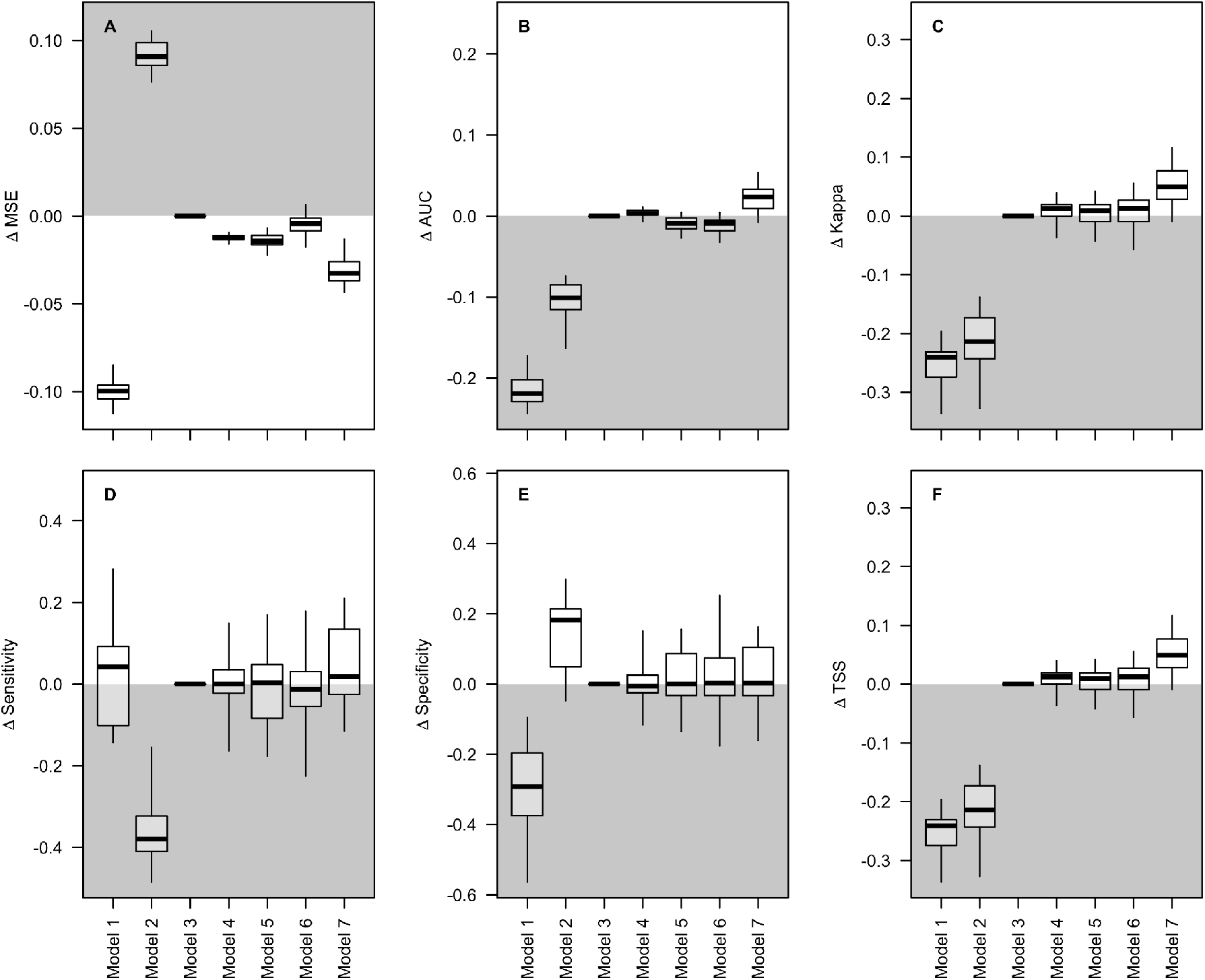
Differences in predictive performance metrics for the encounter rate models 1-7 against balanced and subsampled eBird data from 2017. Metrics are compared to the performance from model 3 and the y-axis values show differences relative to model 3. The white halves of the plots indicate where model performance is *better* than model 3. The grey halves of the plots indicate where model performance is *worse* than model 3. Model 3 uses all the data in a random forest encounter rate model. Model 7 is the random forest encounter rate model using complete checklists, spatial subsampling, effort variable filters, and effort variables as covariates. The validation metrics are calculated for 25 different model runs. For details of models 1-7 see Table 1 and the text. Boxes show the median, the interquartile range, and whisker ends denote the extremes of the distributions.

Models 1 and 2 had substantially worse model performance with both the temporally independent validation and the BBS validation (Figures 3, S2). Their estimates of encounter rate were poorly correlated with those from model 7 (Figure S3), although there are some broad similarities in spatial patterns (Figure 2). These results demonstrate that using presence-only or casual records only is likely to result in poorer ecological inference. Models 3-5 all displayed similar model performance (Figures 3, S2), similar absolute encounter rate (Figure S4), and similar correlations with the predictions from model 7 (Figure S3). All these results suggest that the largest gains in model performance are achieved from using complete checklists (rather than casual records), and including effort variables as covariates.

### Estimating species occupancy

Across models, the estimates of occupancy were less variable than those of encounter rate. The six occupancy models showed relatively consistent spatial patterns (Figure 4), high correlation between estimates (Figure S5), similar absolute estimates of occupancy (Figure S7), and similar model performance at well-monitored sites (Figure S8). Therefore with these training and validation datasets we could not strongly identify improvements with the data processing steps. Despite these apparent similarities, there were large differences in the distributions of estimated detectability within the training data (Figure S6). Therefore using the effort variables did enable the model to account for heterogeneity in detectability, expected to lead to more precise estimates of occupancy.

**Figure 4.**
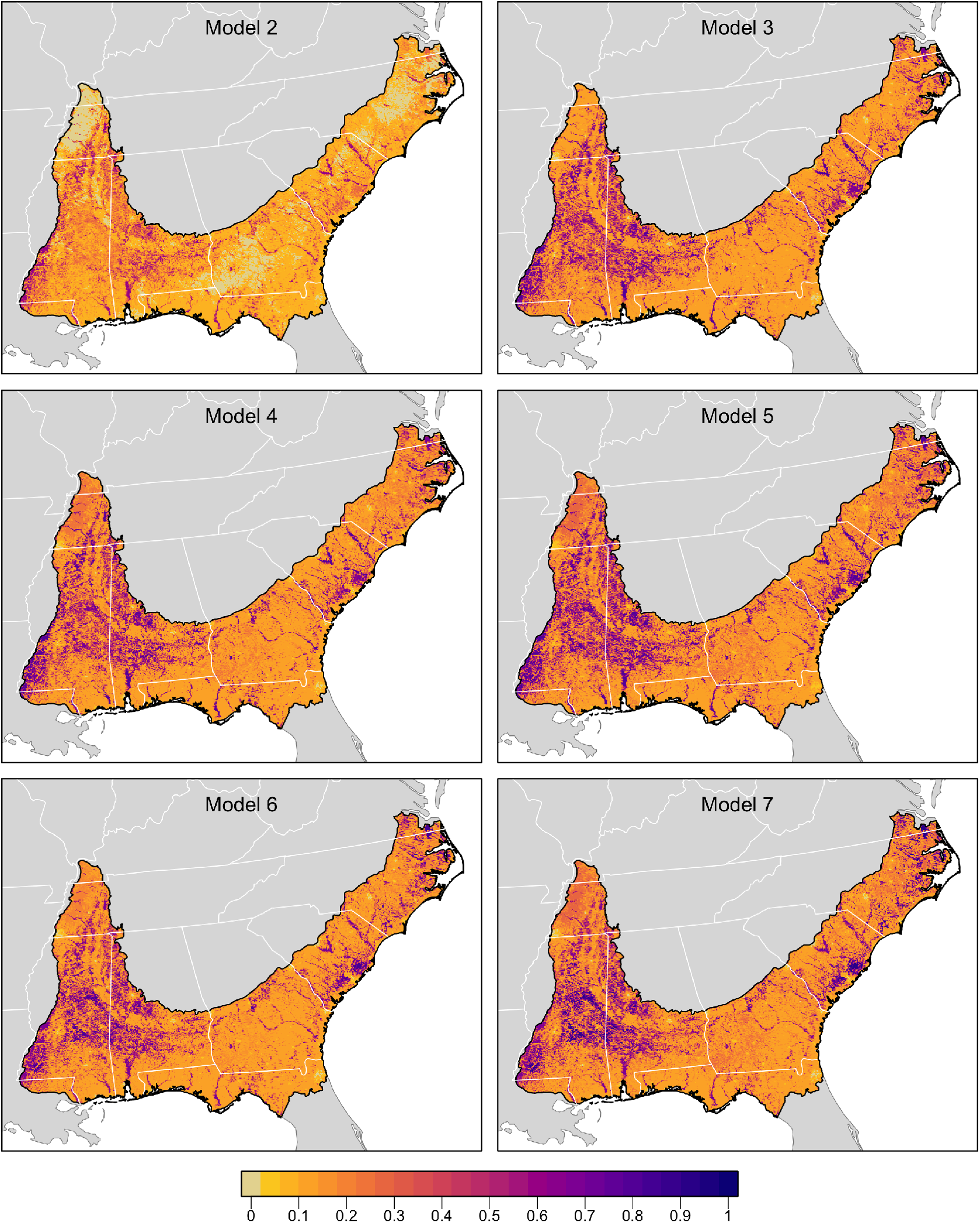
Estimated occupancy of wood thrush across the BCR27 region for occupancy models 2-7 calculated with data processing steps (ii) to (v). The occupancy is the expected probability that cells are occupied by Wood Thrush. Darker colours denote higher estimated occupancy.

### Varying sample size

Model 7 (using all refining data processing steps) was better than model 3 (using all the data without any data processing) (Figure 2). However, the benefits of using model 7 were reduced at smaller sample sizes (Figures 5, S6). This may be because reducing the dataset size by filtering (Figure 1) also has a cost when there are fewer data. However, we find that even with the smallest datasets, there is no disadvantage to using model 7 — it performs equivalent to or better than model 3, across all sample sizes that we tested (Figures 5, S6).

**Figure 5.**
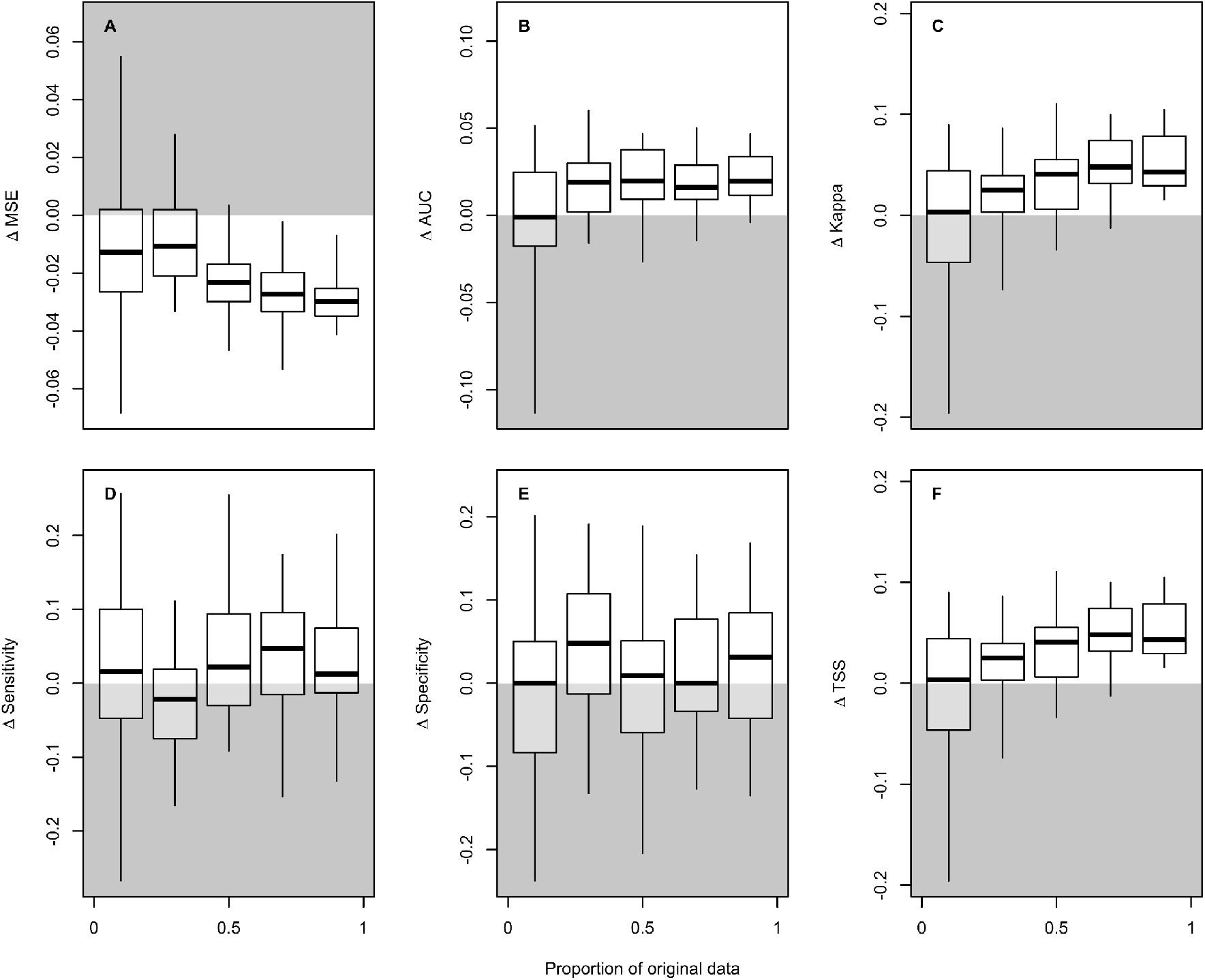
Effect of sample size on differences in predictive performance metrics for the encounter rate models 3 and 7. Differences were computed between the metrics as (model 7 - model 3); the y-axis values show differences relative to model 3. The white halves of the plots indicate where model 7 performance is *better* than model 3. The grey half of the plots indicate where model 7 performance is *worse* than model 3. The test dataset was balanced and subsampled eBird data from 2017. The datasets were random subsampled to 0.75 of the original checklists. Then they were further reduced to a proportion of this dataset: 0.1, 0.3, 0.5, 0.7, 0.9. This process was repeated 25 times to produce 25 paired comparisons of model performance for each dataset size. Each paired comparison between model 3 and model 7 used the same randomly subsampled test and train datasets. See Table 1 for further details of model 3 and model 7. Panels show the following performance metrics: A Mean Squared Error (MSE); B Area Under the Curve (AUC); C Kappa; D sensitivity; E specificity; and F True Skill Statistic (TSS). Boxes show the median, the interquartile range, and the extremes of the distributions.

## Discussion

Citizen science data sets are becoming increasingly valuable research tools for ecology and conservation due to their increasing prevalence (Pocock et al., 2017) and broad spatiotemporal scope (Chandler et al., 2017). For example, eBird data have been used to study phenology, species distributions, population trends, evolution, behaviour, global change, and conservation (Lang, Mann, & Farine, 2018; MacPherson et al., 2018; Mattsson et al., 2018; Mayor et al., 2017; Seeholzer, Claramunt, & Brumfield, 2017). However, citizen science data generally have more errors, assumptions, and biases associated with them, often a result of relatively unconstrained survey design and a highly heterogeneous observation process. Here we demonstrate how thoughtful combinations of data filtering and analysis can remove relatively uninformative data and control for much of the statistical noise in citizen science data.

Spatial subsampling resulted in a greater reduction in dataset size than filtering to remove extreme values of observer effort (Figure 1), although this is dependent on the parameters selected for both of these filtering processes. In our case study, neither of these reductions in sample size led to changes in the accuracy of species distribution models (cf models 3-5), indicating no loss of information was incurred by even the removal of substantial proportions of the data (Figure 1).

Including information on the observation process has been shown to produce more accurate and robust results (Isaac, Van Strien, August, de Zeeuw, & Roy, 2014; Johnston, Fink, Hochachka, & Kelling, 2018). Including effort variables had a larger impact on the accuracy of our encounter rate models than removing records based on extreme observer effort (Figure 2). However, the advantages of effort variables were less apparent with occupancy models, although we recognise that our validation was less strong for the occupancy models. Nonetheless, our results suggest that where these effort data do not exist, occupancy models may be a more robust modelling approach.

We considered the impacts of degrading the data in two separate ways - to detections only (presence only data) and to incomplete checklists only. The data degraded in these two ways led to consistently poorer model performance. There are clear limitations to the ecological insights that can be gained from presence-only data (Aranda & Lobo, 2011; Václavík & Meentemeyer, 2009). As a result, multiple approaches have been suggested for inferring non-detection events when data are stored in a presence-only format (Hill, 2012; van Strien, van Swaay, & Termaat, 2013). Our case study suggests that when complete checklists are available, it is very beneficial to use this information rather than degrade the data quality.

Our general recommendation is that both filtering and modelling variation in observer effort are important analytical tools, however their benefits will vary across datasets and modelling objectives. In our example we find that analysing some complete checklists and using effort variables as covariates made the largest difference to the model quality. However, the raw data, the volume of data, the model type, and the modelling objective will all affect the utility of the data processing steps that we describe. To produce a more comprehensive understanding, To provide a more comprehensive understanding, these filtering and modelling practices should be evaluated in different circumstances and with different datasets.

In conclusion, the data processing steps to refine the data are most relevant to semistructured citizen science projects designed to collect a large quantity of data, and with information describing the observation process (Kelling et al., 2018). There are numerous citizen science programs in the world today, but only a limited number of them collect the information needed to infer absences (Pocock et al., 2017). eBird provides evidence that for birds, information on observer effort and completeness of species lists can be collected whilst maintaining high participation. While we focused on modelling species distributions, many other types of ecological inference will also benefit from these data processing steps. In combination, the approaches outlined here for collecting, processing and modeling citizen science data can inform ways to improve existing and future programmes, while increasing our current capacity to conduct robust analyses using growing volumes of citizen science data.

## Acknowledgements

We thank the thousands of participants in the eBird project. We are grateful to Frank La Sorte, Marshall Iliff, and anonymous editor and reviewers for discussions and comments that improved this manuscript. Funding for this work came from the Wolf Creek Foundation, The Leon Levy Foundation, The Packard Foundation, and the National Science Foundation (grants ITR-0427914, DBI-0542868, IIS-0612031, ABI-1356308, CCF-1522054, ICR-1927646). Funding also came from the 2017-2018 Belmont Forum and BiodivERsA joint call for research proposals, under the BiodivScen ERA-Net COFUND program, with support from the Academy of Finland (AKA), the Swedish Research Council (Formas), the National Research Council of Norway (RCN) and the National Science Foundation (NSF).

## Author contributions

All authors conceived the ideas and designed methodology; AJ analysed the data, based on preliminary analyses conducted alongside WMH, VRG, and OR; AJ, WMH, VRG, ETM, and OR wrote the manuscript. All authors contributed critically to the drafts and gave final approval for publication.

## Data accessibility

In Supporting Information Appendix A4 we provide the subset of data used in this paper with all the code used for the analyses presented here. We will provide a DOI for the github repository A4 on publication.

## Supporting information and appendices

**Appendix A1:** eBird data description

**Appendix A2:** Detailed methods for data processing and model fitting

**Appendix A3:** Fitting species distribution and abundance models with eBird data

**Appendix A4:** Code and data for the analyses in this paper https://github.com/ali-johnston/ebird_analysis_woodthrush

**Figure S1.** Location of checklists for A all training data 2018; B example of training data selecting only complete checklists and then spatially subsampled (data for Model 5); C test data 2017; D test data eBird-entered BBS data 2018.

**Figure S2.** Differences in predictive performance metrics for the encounter rate models 1-7 against BBS data within eBird from 2018, aggregated to BBS routes. Metrics are compared to the performance from model 3, with the y-axis values showing the differences relative to model 3. The white halves of the plots indicate where model performance is *better* than model 3. The grey halves of the plots indicate where model performance is *worse* than model 3. Model 1 is the Maxent model which uses only presences and produces background psuedo-absences. Model 7 is the random forest encounter rate model using complete checklists, spatial subsampling, effort variable filters, and effort variables as covariates. The validation metrics are calculated for 25 different model runs. Further details of models 1-7 are in the text and Table 1. Boxes show the median, the interquartile range, and the extremes of the distributions.

**Figure S3.** Density scatterplots comparing estimates of encounter rate across the BCR27 region for models 1 - 6 against model 7. Brighter colours indicate a higher density of points in that region of the scatterplot. The high densities along the bottom of the figures represent many locations that are predicted to have low encounter rate with models 1-6 and higher encounter rate with model 7. R values show the Pearson correlation coefficient and the associated p-value.

**Figure S4.** Estimated encounter rates across BCR27 for models 1-7. See Table 1, Figure 1, Appendix S2 and the main text for full details of each model.

**Figure S5**. Density scatterplots comparing estimates of occupancy rate across the BCR27 region for models 2 - 6 against model 7. Brighter colours indicate a higher density of points in that region of the scatterplot. R values show the Pearson correlation coefficient and the associated p-value.

**Figure S6**. Estimated occupancy rates across BCR27 for models 2-7. See Table 1, Figure 1, Appendix S2 and the main text for full details of each model.

**Figure S7**. Estimated detectability rates in the training data used for model 7, but predicted from models 2-7. See Table 1, Figure 1, Appendix S2 and the main text for full details of each model.

**Figure S8**. Performance metrics for occupancy models 2-7, created with 2018 data and validated with well-surveyed sites in 2017.

**Figure S9.** Effect of sample size on differences in predictive performance metrics for the encounter rate models 3 and 7. Differences were computed between the metrics as (Model 7 - Model 3). The white half of the plots indicating where Model 7 performance is *better* than Model 3. The grey half of the plots indicating where Model 7 performance is *worse* than Model 3. The test dataset was BBS data within eBird from 2018, aggregated to routes. The datasets were random subsampled to 0.75 of the original checklists. Then they were further reduced to a proportion of this dataset: 0.1, 0.3, 0.5, 0.7, 0.9. This process was repeated 25 times to produce 25 paired comparisons of model performance for each dataset size. Each paired comparison between Model 3 and Model 7 used the same randomly subsampled test and train datasets. See Table 1 for further details of Model 3 and Model 7. Panels show the following performance metrics: A Mean Squared Error (MSE); B Area Under the Curve (AUC); C Kappa; D sensitivity; E specificity; and F True Skill Statistic (TSS). Boxes show the median, the interquartile range, and the extremes of the distributions.

## References

Altwegg, R., & Nichols, J. D. (2019). Occupancy models for citizen-science data. Methods in Ecology and Evolution / British Ecological Society, 10(1), 8–21.

Aranda, S. C., & Lobo, J. M. (2011). How well does presence-only-based species distribution modelling predict assemblage diversity? A case study of the Tenerife flora. Ecography, 34(1), 31–38.

Bird, T. J., Bates, A. E., Lefcheck, J. S., Hill, N. A., Thomson, R. J., Edgar, G. J., … Frusher, S. (2014). Statistical solutions for error and bias in global citizen science datasets. Biological Conservation, 173, 144–154.

Botts, E. A., Erasmus, B. F. N., & Alexander, G. J. (2011). Geographic sampling bias in the South African Frog Atlas Project: implications for conservation planning. Biodiversity and Conservation, 20(1), 119–139.

Casanovas, P., Lynch, H. J., & Fagan, W. F. (2014). Using citizen science to estimate lichen diversity. Biological Conservation, 171, 1–8.

Chandler, M., See, L., Copas, K., Bonde, A. M. Z., López, B. C., Danielsen, F., … Masinde, S. (2017). Contribution of citizen science towards international biodiversity monitoring. Biological Conservation, In press.

Dennis, R. L. H., & Thomas, C. D. (2000). Bias in Butterfly Distribution Maps: The Influence of Hot Spots and Recorder’s Home Range. Journal of Insect Conservation, 4(2), 73–77.

Ellis, M. V., & Taylor, J. E. (2018). Effects of weather, time of day, and survey effort on estimates of species richness in temperate woodlands. Emu - Austral Ornithology, 118(2), 183–192.

Fiske, I., & Chandler, R. (2011). unmarked: An R package for fitting hierarchical models of wildlife occurrence and abundance. Journal of Statistical Software, 43(10), 1–23.

Friedl, M., & Sulla-Menashe, D. (2015). MCD12Q1 MODIS/Terra+ Aqua Land Cover Type Yearly L3 Global 500m SIN Grid V006 [Data set]. NASA EOSDIS Land Processes DAAC. Doi, 10.

Hijmans, R. J., Garrett, K. A., Huaman, Z., Zhang, D. P., Schreuder, M., & Bonierbale, M. (2000). Assessing the geographic representativeness of genebank collections: the case of Bolivian wild potatoes. Conservation Biology: The Journal of the Society for Conservation Biology, 14(6), 1755–1765.

Hill, M. O. (2012). Local frequency as a key to interpreting species occurrence data when recording effort is not known. Methods in Ecology and Evolution / British Ecological Society, 3(1), 195–205.

Howard, E., Aschen, H., & Davis, A. K. (2010). Citizen Science Observations of Monarch Butterfly Overwintering in the Southern United States. Psyche; a Journal of Entomology, 2010. doi: 10.1155/2010/689301

Isaac, N. J. B., Van Strien, A. J., August, T. A., de Zeeuw, M. P., & Roy, D. B. (2014). Statistics for citizen science: extracting signals of change from noisy ecological data. Methods in Ecology and Evolution / British Ecological Society, 5(10), 1052–1060.

Johnston, A., Fink, D., Hochachka, W. M., & Kelling, S. (2018). Estimates of observer expertise improve species distributions from citizen science data. Methods in Ecology and Evolution / British Ecological Society, 9, 88–97.

Kadmon, R., Farber, O., & Danin, A. (2004). Effect of roadside bias on the accuracy of predictive maps produced by bioclimatic models. Ecological Applications: A Publication of the Ecological Society of America, 14(2), 401–413.

Kamp, J., Oppel, S., Heldbjerg, H., Nyegaard, T., & Donald, P. F. (2016). Unstructured citizen science data fail to detect long-term population declines of common birds in Denmark. Diversity and Distributions, 22(10), 1024–1035.

Kelling, S., Johnston, A., Fink, D., Ruiz-Gutierrez, V., Bonney, R., Bonn, A., … Guralnick, R. (2018). Finding the signal in the noise of Citizen Science Observations. bioRxiv. Retrieved from https://www.biorxiv.org/content/early/2018/05/18/326314.abstract

Lang, S. D. J., Mann, R. P., & Farine, D. R. (2018). Temporal activity patterns of predators and prey across broad geographic scales. Behavioral Ecology: Official Journal of the International Society for Behavioral Ecology. doi: 10.1093/beheco/ary133

MacPherson, M. P., Jahn, A. E., Murphy, M. T., Kim, D. H., Cueto, V. R., Tuero, D. T., & Hill, E. D. (2018). Follow the rain? Environmental drivers of Tyrannus migration across the New World. The Auk, 881–894.

Mair, L., & Ruete, A. (2016). Explaining Spatial Variation in the Recording Effort of Citizen Science Data across Multiple Taxa. PloS One, 11(1), e0147796.

Mattsson, B. J., Dubovsky, J. A., Thogmartin, W. E., Bagstad, K. J., Goldstein, J. H., Loomis, J. B., … López-Hoffman, L. (2018). Recreation economics to inform migratory species conservation: Case study of the northern pintail. Journal of Environmental Management, 206, 971–979.

Mayor, S. J., Guralnick, R. P., Tingley, M. W., Otegui, J., Withey, J. C., Elmendorf, S. C., … Schneider, D. C. (2017). Increasing phenological asynchrony between spring green-up and arrival of migratory birds. Scientific Reports, 7(1), 1902.

Miller, D. A. W., Pacifici, K., Sanderlin, J. S., & Reich, B. J. (2019). The recent past and promising future for data integration methods to estimate species’ distributions. Methods in Ecology and Evolution / British Ecological Society, 10(1), 22–37.

NABCI (2000). bird conservation region descriptions: a supplement to the North American Bird Conservation Initiative bird conservation regions map. US NABCI Committee.

Newson, S. E., Evans, H. E., & Gillings, S. (2015). A novel citizen science approach for large-scale standardised monitoring of bat activity and distribution, evaluated in eastern England. Biological Conservation, 191, 38–49.

Oliveira, C. V., Olmos, F., dos Santos-Filho, M., & Bernardo, C. S. S. (2018). Observation of Diurnal Soaring Raptors In Northeastern Brazil Depends On Weather Conditions and Time of Day. The Journal of Raptor Research, 52(1), 56–65.

Pacifici, K., Reich, B. J., Miller, D. A. W., Gardner, B., Stauffer, G., Singh, S., … Collazo, J. A. (2017). Integrating multiple data sources in species distribution modeling: a framework for data fusion. Ecology, 98(3), 840–850.

Phillips, S. (2016). Maxnet: Fitting “maxent” species distribution models with “glmnet.”

Pocock, M. J. O., Tweddle, J. C., Savage, J., Robinson, L. D., & Roy, H. E. (2017). The diversity and evolution of ecological and environmental citizen science. PloS One, 12(4), e0172579.

Pya, N. (2013). scam: Shape constrained additive models.

R Core Team. (2018). R: A Language and Environment for Statistical Computing. Retrieved from https://www.R-project.org/

Sauer, J. R., Pardieck, K. L., Ziolkowski, D. J., Smith, A. C., Hudson, M.-A. R., Rodriguez, V., … Link, W. A. (2017). The first 50 years of the North American Breeding Bird Survey. The Condor, 119(3), 576–593.

Seeholzer, G. F., Claramunt, S., & Brumfield, R. T. (2017). Niche evolution and diversification in a Neotropical radiation of birds (Aves: Furnariidae). Evolution; International Journal of Organic Evolution, 71(3), 702–715.

Strimas-Mackey, M., Miller, E., & Hochachka, W. (2018). auk: eBird Data Extraction and Processing with AWK. R Package Version 0.3.0.

Strimas-Mackey, M., W.M. Hochachka, V. Ruiz-Gutierrez, O.J. Robinson, E.T. Miller, T. Auer, S. Kelling, D. Fink, A. Johnston. 2020. Best Practices for Using eBird Data. Version 1.0. https://cornelllabofornithology.github.io/ebird-best-practices/. Cornell Lab of Ornithology, Ithaca, New York. https://doi.org/10.5281/zenodo.3620739

Sullivan, B. L., Aycrigg, J. L., Barry, J. H., Bonney, R. E., Bruns, N. E., Cooper, C. B., … Kelling, S. (2014). The eBird enterprise: An integrated approach to development and application of citizen science. Biological Conservation, 169, 31–40.

Troudet, J., Grandcolas, P., Blin, A., Vignes-Lebbe, R., & Legendre, F. (2017). Taxonomic bias in biodiversity data and societal preferences. Scientific Reports, 7(1), 9132.

Tulloch, A. I. T., Possingham, H. P., Joseph, L. N., Szabo, J., & Martin, T. G. (2013). Realising the full potential of citizen science monitoring programs. Biological Conservation, 165, 128–138.

Tulloch, A. I. T., & Szabo, J. K. (2012). A behavioural ecology approach to understand volunteer surveying for citizen science datasets. Emu - Austral Ornithology, 112(4), 313–325.

Václavík, T., & Meentemeyer, R. K. (2009). Invasive species distribution modeling (iSDM): Are absence data and dispersal constraints needed to predict actual distributions? Ecological Modelling, 220(23), 3248–3258.

Valavi, R., Elith, J., Lahoz-Monfort, J. J., & Guillera-Arroita, G. (2018). block CV: An r package for generating spatially or environmentally separated folds for k - fold cross-validation of species distribution models. Methods in Ecology and Evolution / British Ecological Society, 67, 617.

van Strien, A. J., van Swaay, C. A. M., & Termaat, T. (2013). Opportunistic citizen science data of animal species produce reliable estimates of distribution trends if analysed with occupancy models. The Journal of Applied Ecology, 50, 1450–1458.

Vianna, G. M. S., Meekan, M. G., Bornovski, T. H., & Meeuwig, J. J. (2014). Acoustic telemetry validates a citizen science approach for monitoring sharks on coral reefs. PloS One, 9(4), e95565.

Wiggins, A., & Crowston, K. (2011). From Conservation to Crowdsourcing: A Typology of Citizen Science. 2011 44th Hawaii International Conference on System Sciences, 1–10.

Wright, M., & Ziegler, A. (2017). ranger: A Fast Implementation of Random Forests for High Dimensional Data in C++ and R. Journal of Statistical Software, Articles, 77(1), 1–17.

