## Appendix A1 for "Analytical guidelines to increase the value of citizen science data: using eBird data to estimate species occurrence"

**SUPPORTING INFORMATION APPENDIX A1**

### eBird Data: Descriptions of data review, data access, and data columns

In our examples we use data from the semi-structured citizen science programme eBird (Sullivan et al., 2014). eBird was created as a comprehensive tool and database for collecting high quality bird observations. The data from eBird are valuable to researchers across the globe, given the project's broad spatial coverage and year-round temporal coverage. The project provides high volumes of data in accessible formats useful to address a broad range of ecological questions. As of January 2019, the database contained nearly 600 million species observations from every country in the world. eBird data have been widely used in scientific research to study phenology (Arab, Courter, & Zelt, 2016; Mayor et al., 2017; Probst, Therrien, Goodrich, & Bildstein, 2017), species distributions (Coxen, Frey, Carleton, & Collins, 2017; MacPherson et al., 2018; Merow, Wilson, & Jetz, 2017), species abundance (Johnston et al. 2015, Robinson et al. 2018), population trends (Horns, Adler, & Şekercioğlu, 2018; Walker & Taylor, 2017), evolution (McEntee, Tobias, Sheard, & Burleigh, 2018; Seeholzer, Claramunt, & Brumfield, 2017; Smith, Seeholzer, Harvey, Cuervo, & Brumfield, 2017), behavior (Freeman & Miller, 2018; Lang, Mann, & Farine, 2018), global change (La Sorte et al., 2017; La Sorte, Fink, & Johnston, 2018), and conservation (Belaire, Kreakie, Keitt, & Minor, 2014; Jarvis, Bollard Breen, Krägeloh, & Billington, 2015; Mattsson et al., 2018).

eBird data are subject to a number of quality controls before they are made available. Observations of unexpected species or unusually high numbers of individuals are automatically flagged during the data-entry process, based on human-generated 'filters'. There are three sequential ways these flagged records are processed: 1) Users are notified of the unusual nature of the record when entering the data, providing them an opportunity to check for errors or reassess their identification. 2) Participants are requested to provide additional information for their observation, 3) The unusual observations are assessed by expert volunteer regional reviewers, who validate the observation based on the additional information. Unusual observations that pass through all of these processes are made accessible in the eBird public database, otherwise they remain hidden from public view. The species and numbers of individuals that are used to identify unusual observations are spatiotemporally defined, because 'unusual' varies by both location and season. In some countries species previously unreported in the country are flagged, whereas in areas with much data, flagged species can be those unknown from specific locations, or at specific times of year. As a corollary, records that meet expectations enter the database unaltered and without human vetting, meaning the degree to which 'expected' species are erroneously reported (i.e., false positives) is currently unknown.

For some species the number of false positives may be problematic, particularly if they are rare species that are similar in appearance to more common species.

There are two key aspects of eBird data that enable robust ecological conclusions. Firstly, some checklists are recorded as 'complete checklists' where all birds detected and identified are recorded on the checklist and thus allowing for non-detections to be inferred. Secondly, eBird collects information on the effort expended during the observation process. This aspect provides important information to describe the observation process and qualifies eBird as a semi-structured citizen science project (Kelling et al., 2018).

**eBird data access**

The eBird Basic Dataset (EBD) is a data product created by the Cornell Lab of Ornithology both for internal use and for public distribution, and is a "flat file" summary of the information contained in the project's underlying relational database. The EBD is global in extent and updated monthly (www.ebird.org/science/download-ebird-data-products). Both the entire global EBD and coarsely defined subsets (based on political regions, species, and date ranges) can be downloaded. While access to the EBD data product requires completion of a data-access request, this access is unrestricted for non-commercial uses. While much of the information contained in the EBD is also archived with the Global Biodiversity Information Facility (GBIF), we discourage the use of eBird data as obtained from GBIF. The data in GBIF are reduced to presence-only format and also lose information on the effort associated with each observation. As we demonstrate in this paper, both of these losses of information have a detrimental effect on the accuracy of models produced with these data.

Associated with the EBD data product is an R package, *auk* (Strimas-Mackey, Miller, & Hochachka, 2017) that is designed to facilitate three data-preparation tasks: (1) extraction of data for specific species, regions, and ranges of dates; (2) filtering of these data in order to minimize the range of observer effort contained within the data; and (3) converting the data in the EBD, which are stored in a detection-only format, into a "detection/non-detection" or "count/non-detection" format. We recommend that the raw EBD data are always converted into "detection/non-detection" form before use. The supporting information A4 bookdown document presents an example of the use of the *auk* package.

**Important eBird concepts**

There are several features of the data within eBird that need to be accounted for by anyone wishing to use these data, and which do not fit conveniently within the scope of the generic processes of filtering data and modelling variation in detection.

*Shared checklists*

Bird watching is often a social activity, and when bird watching in groups, each individual will typically want to have their own records of the species that they saw. To accommodate this situation, the eBird database contains the concept of the ‘shared checklist’. With a shared checklist, a single observer records species and submits the checklist into the eBird database. The observer then shares this list with other members of their group. Within the database, this process of sharing a checklist results in the creation of an almost identical checklist except for the fact that this new copy is associated with a different observer (i.e. the *OBSERVER ID* is different). All copies of a shared checklist are connected by a unique *GROUP ID* value within the database. Once a checklist is shared, each copy’s owner can modify any aspect of the contents of their version of the checklist, so the checklists can vary between observers, even though they are from the same original checklist. To the best of our knowledge almost all modifications are the additions or removals of only a few species, or small changes in the numbers of individual birds observed, which reflects the fact that not all members of a bird watching group will see all individuals of all species.

Because of the redundancy of information across shared checklists, any analysts will most often want to collapse down the information in each group of shared checklists into a single observation. The *auk* R package by default will collapse groups of shared lists into a single entity by retaining only the first-created checklist within a shared group when reading the data into R using the *read_ebd* function. Other methods of collapsing information across shared checklists are possible, albeit with greater computational expense, which some analysts may find appropriate.

*eBird taxonomy*

eBird uses its own taxonomic system, which very closely maps onto other global avian taxonomies and nomenclatures. We emphasise that for any eBird analysis, researchers should consider the taxonomy of their study organisms, and whether eBird’s taxonomy is appropriate for their intended use of the data. In many cases taxonomic differences can be dealt with by querying specific subspecies, or multiple species, and splitting and lumping after the query as necessary (“The eBird Taxonomy,” n.d.). See the eBird website for further details: https://help.ebird.org/customer/portal/articles/1006825-the-ebird-taxonomy.

The eBird taxonomy is updated annually after expert review, and changed to reflect new data or concensus. A notable benefit of these annual, system-wide updates is that splits and lumps of species are propagated across the entire historical eBird database. When necessary, expert reviewers manually sort through observations and assign records to current taxonomic hypotheses to the extent possible.

The eBird taxonomy allows for taxonomic units other than species, including subspecies. Some observers record birds' identities at these finer resolutions and this information is stored in the database. For many or most purposes, these finer-resolution identities should be translated to the species level. The *read_ebd* function of *auk* by default conducts this translation, but this behaviour can be disabled if information is required at the subspecies level.

**Controlling bias and variance with eBird data**

Similar to many citizen science projects, eBird requires no specific structure regarding the location, date, time, or protocol for each observation event. However, unlike some other projects, ancillary data are collected that allow *post hoc* control for variation in the observation process (Kelling et al., 2018), This control occurs by filtering of the data to limit the range of variation present in the data to be analysed, and by the inclusion of predictor variables in models to account for sources of variation. Below, we describe how the information contained in the EBD can be used to address the issues that the main paper describes as the major sources of potential bias and imprecision with eBird data and many citizen science projects.

*Spatial precision*

For all eBird checklists the spatial location is provided as a single latitude-longitude point. However, there are two main reasons why this location may not be a precise representation of the locations at which birds were detected. Firstly, for traveling checklists the point represents only one location on the route travelled. Secondly, a checklist location may be assigned to a 'hotspot' (a location that is expected to be visited by many bird watchers). However, participants that assign their checklist to a hotspot may not be at the precise hotspot location, even for stationary checklists. It is therefore not appropriate to align the eBird locations with very precise and fine-resolution measures of environmental covariates. For these two reasons, in many of our analyses we match the checklist with environmental covariates within approximately a 3 km $\times$ 3 km square centred on the checklist point. For example, in this study we use a 2.5 km $\times$ 2.5 km square. There is some trade-off between the size of this square and the maximum distance of traveling count used. We have found that for travelling checklists up to 5km in length, the vast majority of bird-watching activity took place within the 3km grid square centred on the locations designated for these travelling checklists. Note that even when using stationary checklists we suggest caution with the interpretation of spatial precision.

*Spatial sampling bias*

Locations of eBird checklists reflect the natural spatial bias in the locations selected by bird watchers. For example, there are preferences of bird watchers to engage in their hobby in convenient locations (i.e. often close to their home), as well as areas that are popular because they have high avian diversity. For example, most participants in citizen science surveys sample near their homes (Luck, Ricketts, Daily, & Imhoff, 2004), easily accessible areas such as roadsides (Kadmon, Farber, & Danin, 2004), or in areas and habitats of known high biodiversity (Prendergast, Wood, Lawton, & Eversham, 1993). A simple method for reducing spatial bias is to create a grid over the region of interest, and sample a fixed number of checklists from within each grid cell. Care should be taken when selecting the size of the grid cell sizes, as this parameter controls the tradeoff between bias control and sample size, which affects model variance. Selecting a small size can cause the grid-sampled data to retain strong spatial bias because there will be many small grid cells with no observations and regions with high densities of observations will still have a higher density of data after spatial subsampling. However, selecting a large size of grid cell size can substantially reduce the sample size. Selecting a high number of checklists per grid cell to combat this will lead to data that retain spatial bias. Our experience suggests that the spatial precision at which spatially varying predictors are described is a useful starting point in deciding upon the sizes of grid cells; this is why we have used a grid with 5km distance between cell centres in the examples presented with this paper. It should also be noted that this spatial subsampling reduces the larger-scale spatial bias between grid cells, resulting from proximity to where participants live, but may not reduce the finer-scale spatial bias within grid cells resulting from preferential selection of certain habitat types. Finally, for species that are not common in the data, subsampling can create a dataset with very few positive observations. Although this doesn't change the proportion of positives in the data, if the absolute number gets too low, then methods to address class imbalance may be appropriate (see Appendix A3).

*Temporal bias*

The number of checklists submitted to eBird increases every year, with exponential increases typically found in annual growth for a given region, as eBird is adopted. As a result, information from the most recent years can mask any patterns in data from earlier years. For example, changes in habitat association through time, such as increased colonisation of urban areas through time (Evans, Hatchwell, Parnell, & Gaston, 2010), can remain undetected unless appropriate steps are taken to account for the potential masking effects of increases in sample sizes through time. As with other aspects of addressing potential biases in analyses of citizen science data, a combination of data filtering and use of appropriate predictor variables in models should be used to account for temporal bias. Filtering for temporal bias is analogous to filtering for spatial bias (see *Spatial bias*, above): selecting a random (or other appropriate) subset of data from each year so that there is an approximately equal representation of data from all years. Following filtering the inclusion of year as a predictor variable (main effect, or in interactions) should be able to describe patterns of change across years.

The same general considerations for dealing with increases in the abundance of data across years are also applicable for handling systematic variation in the volume of data within an individual year. Bird watchers, on average, are not equally enthusiastic about their hobby throughout the year. For example, we see clear indications in North America of greater bird watching effort in the late winter to early spring and a distinct drop in bird watchers' activity in the middle of summer (see Figure 1 of (Sullivan et al., 2014)). Seasonal balancing of data should be considered when research interests involve describing cross-season variation in behaviour or local (i.e. habitat-related) distribution.

**References**

Arab, A., Courter, J. R., & Zelt, J. (2016). A spatio-temporal comparison of avian migration phenology using Citizen Science data. *Spatial Statistics*, *18*, 234–245.

Belaire, J. A., Kreakie, B. J., Keitt, T., & Minor, E. (2014). Predicting and Mapping Potential Whooping Crane Stopover Habitat to Guide Site Selection for Wind Energy Projects: Whooping Cranes and Wind Farms. *Conservation Biology: The Journal of the Society for Conservation Biology*, *28*(2), 541–550.

Coxen, C. L., Frey, J. K., Carleton, S. A., & Collins, D. P. (2017). Species distribution models for a migratory bird based on citizen science and satellite tracking data. *Global Ecology and Conservation*, *11*, 298–311.

Evans, K. L., Hatchwell, B. J., Parnell, M., & Gaston, K. J. (2010). A conceptual framework for the colonisation of urban areas: the blackbird Turdus merula as a case study. *Biological Reviews of the Cambridge Philosophical Society*, *85*(3), 643–667.

Freeman, B. G., & Miller, E. T. (2018). Why do crows attack ravens? The roles of predation threat, resource competition, and social behavior. *The Auk*, *135*(4), 857–867.

Horns, J. J., Adler, F. R., & Şekercioğlu, Ç. H. (2018). Using opportunistic citizen science data to estimate avian population trends. *Biological Conservation*, *221*, 151–159.

Jarvis, R. M., Bollard Breen, B., Krägeloh, C. U., & Billington, D. R. (2015). Citizen science and the power of public participation in marine spatial planning. *Marine Policy*, *57*, 21–26.

Kadmon, R., Farber, O., & Danin, A. (2004). EFFECT OF ROADSIDE BIAS ON THE ACCURACY OF PREDICTIVE MAPS PRODUCED BY BIOCLIMATIC MODELS.

*Ecological Applications: A Publication of the Ecological Society of America*, *14*(2), 401–413.

Kelling, S., Johnston, A., Fink, D., Ruiz-Gutierrez, V., Bonney, R., Bonn, A., … Guralnick, R. (2018). Finding the signal in the noise of Citizen Science Observations. *bioRxiv*. Retrieved from https://www.biorxiv.org/content/early/2018/05/18/326314.abstract

Lang, S. D. J., Mann, R. P., & Farine, D. R. (2018). Temporal activity patterns of predators and prey across broad geographic scales. *Behavioral Ecology: Official Journal of the International Society for Behavioral Ecology*. doi: 10.1093/beheco/ary133

La Sorte, F. A., Fink, D., Blancher, P. J., Rodewald, A. D., Ruiz-Gutierrez, V., Rosenberg, K. V., … Kelling, S. (2017). Global change and the distributional dynamics of migratory bird populations wintering in Central America. *Global Change Biology*, *23*(12), 5284–5296.

La Sorte, F. A., Fink, D., & Johnston, A. (2018). Seasonal associations with novel climates for North American migratory bird populations. *Ecology Letters*, *21*(6), 845–856.

Luck, G. W., Ricketts, T. H., Daily, G. C., & Imhoff, M. (2004). Alleviating spatial conflict between people and biodiversity. *Proceedings of the National Academy of Sciences of the United States of America*, *101*(1), 182–186.

MacPherson, M. P., Jahn, A. E., Murphy, M. T., Kim, D. H., Cueto, V. R., Tuero, D. T., & Hill, E. D. (2018). Follow the rain? Environmental drivers of Tyrannus migration across the New World. *The Auk*, 881–894.

Mattsson, B. J., Dubovsky, J. A., Thogmartin, W. E., Bagstad, K. J., Goldstein, J. H., Loomis, J. B., … López-Hoffman, L. (2018). Recreation economics to inform migratory species conservation: Case study of the northern pintail. *Journal of Environmental Management*, *206*, 971–979.

Mayor, S. J., Guralnick, R. P., Tingley, M. W., Otegui, J., Withey, J. C., Elmendorf, S. C., … Schneider, D. C. (2017). Increasing phenological asynchrony between spring green-up and arrival of migratory birds. *Scientific Reports*, *7*(1), 1902.

McEntee, J. P., Tobias, J. A., Sheard, C., & Burleigh, J. G. (2018). Tempo and timing of ecological trait divergence in bird speciation. *Nature Ecology & Evolution*, *2*(7), 1120–1127.

Merow, C., Wilson, A. M., & Jetz, W. (2017). Integrating occurrence data and expert maps for improved species range predictions: Expert maps & point process models. *Global Ecology and Biogeography: A Journal of Macroecology*, *26*(2), 243–258.

Prendergast, J. R., Wood, S. N., Lawton, J. H., & Eversham, B. C. (1993). Correcting for Variation in Recording Effort in Analyses of Diversity Hotspots. *Biodiversity Letters*, *1*(2), 39–53.

Probst, J. C., Therrien, J.-F., Goodrich, L. J., & Bildstein, K. L. (2017). Increase in Numbers and Potential Phenological Adjustment of Ruby-throated Hummingbirds (Archilochus colubris) during Autumn Migration at Hawk Mountain Sanctuary, Eastern Pennsylvania, 1990--2014. *The Wilson Journal of Ornithology*, *129*(2), 360–364.

Seeholzer, G. F., Claramunt, S., & Brumfield, R. T. (2017). Niche evolution and diversification in a Neotropical radiation of birds (Aves: Furnariidae). *Evolution; International Journal of*

*Organic Evolution*, *71*(3), 702–715.

Smith, B. T., Seeholzer, G. F., Harvey, M. G., Cuervo, A. M., & Brumfield, R. T. (2017). A latitudinal phylogeographic diversity gradient in birds. *PLoS Biology*, *15*(4), e2001073.

Strimas-Mackey, M., Miller, E., & Hochachka, W. (2017). auk: eBird Data Extraction and Processing with AWK. *R Package Version*.

Sullivan, B. L., Aycrigg, J. L., Barry, J. H., Bonney, R. E., Bruns, N. E., Cooper, C. B., … Kelling, S. (2014). The eBird enterprise: An integrated approach to development and application of citizen science. *Biological Conservation*, *169*, 31–40.

The eBird Taxonomy. (n.d.). Retrieved November 5, 2018, from eBird website: https://help.ebird.org/customer/portal/articles/1006825-the-ebird-taxonomy

Walker, J., & Taylor, P. (2017). Using eBird data to model population change of migratory bird species. *Avian Conservation and Ecology/Ecologie et Conservation Des Oiseaux*, *12*(1). Retrieved from https://www.ace-eco.org/vol12/iss1/art4/
