## Appendix A2 for "Analytical guidelines to increase the value of citizen science data: using eBird data to estimate species occurrence"

**SUPPORTING INFORMATION APPENDIX A2**

### Data processing and model descriptions

This appendix lists in detail the methods used for data processing and model fitting for the analyses in this paper. The R code used for data processing and fitting the models in this paper is available in supporting information A4: https://github.com/ali-johnston/ebird_analysis_woodthrush.

We created seven encounter rate models and six occupancy models based on five data processing steps and different model structures (Table 1). Two of the data processing steps will degrade the data, and three of the data processing steps refine the data. We used these five data processing steps to create different datasets for the various models. Here we outline the data processing steps and then we describe the seven encounter rate models and six occupancy models.

#### Data processing steps

*i) Select detections only*

Here we select locations with detections, removing all non-detections from the data. This provides information about where the species was recorded, but not about where it was not recorded. Many analyses of eBird data reduce the data to detections only.

*ii) Select incomplete checklists only*

Many citizen science projects do not collect information on whether a checklist is complete, so to replicate this we selected only incomplete checklists. Here we infered non-detection, creating a binary 0/1 column with 0 for checklists without non-detections, a process known as “zero-filling”. However with incomplete checklist non-detection is actually “not recorded”, which can arise a number of reasons. The “zeros” produced from incomplete checklists are a combination of ecological processes, observation processes, and species bias. We conducted the zero-filling process with the R package *auk* (Strimas-Mackey, Miller, & Hochachka, 2017). However, it is not recommended to zero-fill incomplete checklists and the *auk* package by default will not do this.

*iii) Select complete checklists only*

The second step was to reduce the full dataset to contain only those data from complete checklists (i.e., the observer indicated that they reported all species that they detected and identified). This step creates a more standardised dataset; a non-detection on a complete checklist is always a non-detection, so the zero-filling process described above is more justified for complete checklists. We used the indicator column *all_species_recorded* to select only complete checklists.

*iv) Spatial subsampling.*

The spatial bias in unstructured or semi-structured citizen science data gives rise to considerably more data from some habitats and locations. Without spatial subsampling this can lead to the estimated relationships between species and environment dominated by places and habitats represented by much more data. These data were also temporally subsetted by randomly selecting a single checklist from within each calendar week. Spatial subsampling reduces the influence of places with many checklists. However, it cannot create data where there are no checklists. So spatial subsampling mitigates, but does not remove, spatial bias.

In order to conduct spatial subsampling, we defined an equal area hexagonal grid across the region, with roughly 5 km between the centres of adjacent hexagons, using the R package *dggridR* (Barnes et al. 2017). We randomly selected one checklist from each hexagon from each week during May 15 - June 30 in 2018.

*v) Effort variables*

We know from experience that variation in the eBird observation process is often the most important source of variation in the likelihood of recording a species. There are five important effort variables associated with each checklist: the time observations started, date, duration of observation process, distance travelled, protocol (e.g., ‘stationary’ and ’travelling’ counts), and the number of observers.

We filtered by these variables to remove more extreme checklists to create a more standardised range of effort within the final dataset. We kept only checklists that used a ‘Stationary’ or ‘Traveling’ protocol. We removed checklists with: durations > 5 hours; distance > 5km; and number of observers > 10). All of these filters together removed only 8% of complete checklists, so effectively this step removes those records for which the effects of variation in effort could not be estimated with high precision. In supporting information A2 we provide further details of the EBD variables that can be used to model the observation process.

**Environmental variables**

As environmental covariates we used land cover data derived from MODIS product MCD12Q1 v006 (Friedl & Sulla-Menashe, 2015). We estimated the land cover associated with each checklist as the proportion of each land cover category in a 2.5 km $\times$ 2.5 km square surrounding the checklist location in the year the observations were made. We included the proportions of each land cover type in the MCD12Q1 v006 classification.

The landcover types we used were:

Class 0: Water bodies

Class 1: Evergreen Needleleaf Forests

Class 2: Evergreen Broadleaf Forests

Class 3: Deciduous Needleleaf Forests

Class 4: Deciduous Broadleaf Forests

Class 5: Mixed Forests

Class 6: Closed Shrublands

Class 7: Open Shrublands

Class 8: Woody Savannas

Class 10: Grasslands

Class 11: Permanent Wetlands

Class 12: Croplands

Class 13: Urban and Built-up Lands

Class 14: Cropland/Natural Vegetation Mosaics

Class 15: Non-Vegetated Lands

**Encounter rate models**

The seven models in the encounter rate set were constructed as follows:

*Encounter rate model 1*

This model used as input all records of detections from the eBird dataset, so we ran a typical ‘presence-only’ Maxent model. We randomly generated 10000 background pseudo-absences and used the *maxnet* package in R (Phillips, 2016). Environmental variables included the 16 landcover covariates. No effort covariates were included and the model was not calibrated, because calibration is not possible with presence only data.

The outputs from the Maxent model represent relative rankings of the sites for encounter rate, but they do not relate to actual encounter rates. The scale is therefore effectively irrelevant.

*Encounter rate model 2*

This model was fitted to checklists from the original training data (subsampled to 0.75 of the original number), but filtered to **incomplete** checklists. The data used to fit Model 2 are our attempt to approximate the type of data that is collected in many biodiversity reporting schemes, for which there is no attempt or incentive to have observers recording the observations of all of the species that they observed. We fitted a random forest model with the R package *ranger* (Wright & Ziegler, 2017). The response was the binary detection/non-detection of a wood thrush on a checklist. Environmental covariates were the 16 landcover variables. There were no effort covariates. We fitted a probability forest to estimate the probability of encounter of wood thrush on a checklist. We fitted 1000 trees and used the square root of the number of covariates (4) to split at in each node (James, Witten, Hastie, & Tibshirani, 2013).

After fitting the model it can be important to calibrate the resulting predictions results (Dormann, 2020). To calibrate, we predicted probability of occurrence at each of the training checklists using the random forest described above. We then modelled the observed detection/non-detection against the predicted probability. To fit this relationship we used a Generalized Additive Model (GAM) that was constrained to be monotonically increasing with the R packages *scam (Pya, 2013)* and *mgcv (Wood, 2006)*. For the smooth we used 4 degrees of freedom and a gamma penalty of 1.4 (Wood, 2006).

For predicting, we first predicted encounter rate with the random forest, then calibrated these predictions with the constrained GAM. The final prediction was a combination of these two models. The estimated encounter rate was: the estimated encounter by an average eBirder on an average incomplete checklist.

*Encounter rate model 3*

This model was fitted to all the checklists from the original training data (subsampled to 0.75 of the original number). As with model 2, we fitted the data with used a probability random forest with 16 landcover covariates, and a constrained GAM for calibration. As above, the estimated encounter rate was: the estimated encounter by an average eBirder on an average checklist.

*Encounter rate model 4*

This model used the checklists from model 3, but filtered these to only the complete checklists with step (iii) above. As with model 3, we fitted the data with a probability random forest with 16 landcover covariates, and a constrained GAM for calibration. As above, the estimated encounter rate was the estimated encounter by an average eBirder on an average complete checklist.

*Encounter rate model 5*

This model used the complete checklists that were zero-filled (from Model 4), but the data were spatiotemporally subsetted as described in step (iv) above. This significantly reduced the number of checklists available to less than half those available for model 4 (Figure 1). As with model 4, we used a probability random forest with 16 landcover covariates, and a constrained GAM for calibration. Also as above, the estimated encounter rate was: the estimated encounter by an average eBirder on an average complete checklist.

*Encounter rate model 6*

This model uses the zero-filled, complete checklists, spatially subsampled data from Model 5. But additionally it uses effort filters to remove some of the checklists with more extreme effort at the tails of the effort distributions, described in step (v) above. As with model 5, we used a random forest with 16 landcover covariates and a constrained GAM for calibration. Again as above, the estimated encounter rate was: the estimated encounter by an average eBirder on an average complete checklist.

*Encounter rate model 7*

This model uses the same data as model 6, but additionally includes effort variables as covariates. The effort variables included as covariates were: day of year (as a continuous variable between May 15 and June 30), the time observations started (as a decimal number between 0 and 24), the duration of the checklist (in minutes), the number of observers, whether the observer was stationary or travelling, and if travelling the distance travelled (in km). As above we used a random forest to estimate encounter rate and a constrained GAM for calibration.

Producing predictions was a little more involved for this model, as we also needed to provide values for the effort variables in the predictor set. For most of these we used the following standardised values: June 15, 1 observer, 1 km travelling count, 1 hour duration. However for time of day we optimised it to the time of day at which encounter was highest. We expect the partial effect of time of day to mostly describe variation in detectability due to time of day (affected by light, noise, bird behaviour). We therefore used the marginal relationship between time and encounter (partial dependence) to estimate the time at which detectability was highest, as it makes sense to predict for a high detectability time of day. To estimate the time of day with maximum encounter rate we first searched for 10 min periods during 24 hours with at least 20 checklists or at least 0.05% of checklists. This ensured that we were only using a partial dependence at times of day when there was good data to inform it. We then searched through these times to identify the time with the highest partial dependence, using the *partial_dependence* function in *ranger* to generate the partial dependence. As this was a relatively time-consuming step, we first used a coarse search to narrow down the time, and then a fine-scale search by 10 minute periods. This process meant that the estimated encounter rate was: the estimated encounter by an average eBirder travelling 1km for 1 hour on June 15, at the optimal time of day for detecting the species.

**Occupancy models**

The six models in the occupancy set were constructed as follows.  It is not possible to fit an occupancy model to detection-only data, and hence no Model 1 exists. They are named Model 2 - Model 6 to keep them aligned with the encounter rate models.

*Occupancy model 2*

This model was fitted to checklists from the original training data, but filtered to only **incomplete** checklists. The data used to fit Model 2 are our attempt to approximate the type of data that is collected in many biodiversity reporting schemes, for which there is no attempt or incentive to have observers recording the observations of all of the species that they observed. The checklists were then combined into ‘sites’ that have the same location information. Sites with at least 2 visits (checklists) were retained and those with more than 10 visits were randomly reduced to keep 10 random visits (checklists).

A single season occupancy model was run using the R package *unmarked (Fiske & Chandler, 2011)*. Four environmental covariates were included in the occupancy submodel that were *a priori* thought to be associated with wood thrush occupancy, based on ecological knowledge. These were based on MODIS satellite data, classified and summarised with 3 km squares surrounding a checklist. The environmental covariates were: deciduous and mixed forest (thought to be positively associated with woodthrush occupancy) and cropland and urban (thought to be negatively associated with woodthrush occupancy). Day of year was included as a covariate of detectability. In reality, both occupancy and detectability will vary with day of year. Including day of year only in the detectability component will mean that the predicted occupancy will be for the day during the data time period with the highest occupancy. Occupancy rate was predicted in each 3km grid cell in the study area. The estimated occupancy rate was: the estimated occupancy of wood thrush in the regions surveyed by people doing checklists at a given location.

*Occupancy model 3*

This model was fitted to all the checklists from the original training data. As with occupancy model 2, we fitted the data with a single season occupancy model. As above, the estimated occupancy rate was: the estimated occupancy of wood thrush in the regions surveyed by people doing checklists at a given location.

*Occupancy model 4*

This model used the checklists from model 3, but filtered these to only the complete checklists with step (iii) above. As with occupancy model 3, we fitted the data with a single season occupancy model. As above, the estimated occupancy rate was: the estimated occupancy of wood thrush in the regions surveyed by people doing checklists at a given location.

*Occupancy model 5*

This model used the complete checklists that were zero-filled (from model 4), but the data were additionally spatiotemporally subsetted as described in step (iv) above. This occurred after aggregating the data into sites, each with 2-10 visits. A maximum of one ‘site’ was randomly selected within each 5km hexagon across the study area. As with occupancy model 4, we fitted the data with a single season occupancy model with four environmental covariates. As above, the estimated occupancy rate was: the estimated occupancy of wood thrush in the regions surveyed by people doing checklists at a given location.

*Occupancy model 6*

This model uses the zero-filled, complete checklists, from model 4. But additionally it uses effort filters to remove some of the checklists with more extreme effort at the tails of the effort distributions, described in step (v) above. The effort filters were applied to the checklists and then they were combined into 'sites' with 2-10 visits. After this step, spatial subsampling was conducted as with model 5. As with occupancy model 5, we fitted the data with a single season occupancy model with four environmental covariates. As above, the estimated occupancy rate was: the estimated occupancy of wood thrush in the regions surveyed by people doing checklists at a given location.

*Occupancy model 7*

This model uses the same data as occupancy model 6, but additionally includes effort variables as covariates. The effort variables included as covariates were: day of year (as a continuous variable between May 15 and June 30), the time observations started (as a decimal number between 0 and 24), the duration of the checklist (in minutes), the number of observers, whether the observer was stationary or travelling, and if travelling the distance travelled (in km). As with occupancy model 6, we fitted the data with a single season occupancy model with four environmental covariates, but additionally these six detectability covariates. As above, the estimated occupancy rate was: the estimated occupancy of wood thrush in the regions surveyed by people doing checklists at a given location. Unlike with encounter rate model 7, we did not need to define effort covariates for the predictions, because the model separates the occupancy and detectability processes. So we can predict occupancy without requiring detectability covariate values.

**Model validation**

For our assessments we wanted to evaluate models based on their ability to produce accurate predictions across the entire spatial extent of our study region. For this purpose, it is appropriate to create a spatially balanced set of test data that evenly represents the study region. We also wanted to assess the ability of the models to discriminate between places of species presence and absence, so we created a balanced validation dataset with equal detections and non-detections. With the encounter rate models, the best way to do this is by assessing the ability of the models to discriminate between detections and non-detections in the validation dataset. We selected two different validation datasets for the encounter rate models (temporal independence and structural independence), and one validation dataset for the occupancy models (with high detectability and temporal independence).

*Encounter rate model validation 1*

The first encounter rate validation dataset was based on temporally independent data. We used the non-BBS data from eBird within the period May 15 - June 30 2017. To produce variation in the test set, for each of the 25 runs, we randomly selected 75% of the validation data. We then selected only complete checklists and those without extreme effort. To assess the ability of the models to discriminate, we wanted to balance the number of detections and non-detections. To assess the ability of the models to predict across the study area, we wanted to spatially subsample the data. A raw spatial subsample of the data would result in few detections, making it challenging to balance detections and non-detections. Therefore, we separated the detections and non-detections and ran a spatial subsample on each set separately. As with the training data we randomly selected 1 checklist from each 5km hexagon within the study area. We then further randomly reduced the set of spatially subsampled non-detections so the number equalled the number of spatially subsampled detections. This process is shown schematically in Figure 1 of the main paper.

*Encounter rate model validation 2*

Our second validation data set is independent of the data used for fitting encounter-rate models by coming from a highly structured, annual bird survey, the North American Breeding Bird Survey (BBS) (Sauer et al., 2017). This is the most widely used source of information on population trends of breeding birds in North America (Sauer & Link, 2011). The BBS data collection process is highly structured, with pre-selected sampling locations, a narrowly defined range of dates that coincide with typical breeding seasons and with the time of day and durations fixed. For some species, such as wood thrush, this more structured dataset is a good validation set. However, BBS routes are 25 miles long and contain 50 evenly spaced stops with 3 minute point counts. The locations of the point counts are not available, however for some BBS routes we were able to acess these from eBird.

Some BBS surveyors input their BBS surveys into eBird. We identified these as series of several 3 minute stationary counts, by the same individual on the same day, each of which was within a temporal and spatial tolerance from the adjacent stops in the sequence (20 mins, 1.5km). The checklists identified as BBS point counts were not very sensitive to changes to these values. We removed these BBS counts from the eBird training data and the 2017 validation data described above. We selected the BBS counts from 2018 to use as the second validation set. We predicted the estimated encounter rate at each point count. Then we aggregated these to the route level - calculating the probability of wood thrush encounter at some point in the route as 1- sum(1-$p_i$) where $p_i$ is the estimated probability of wood thrush at each 3 min stationary point count. We compared this to whether wood thrush was observed at any point in the route, which resulted in 38 routes in 2018 within our region that we could use for validation. We calculated this aggregated measure across the route, because BBS data are mostly analysed at the route level.

*Occupancy model validation*

Occupancy models, due to their nature as two linked logistic regressions (describing detection and occupancy processes), are problematic for cross validation, and no generally accepted method currently exists. To specifically validate the occupancy part of the model, we wanted to identify locations with high detectability. We took the validation data from 2017 before it had been spatially subsampled. We predicted detectability on each checklist and then created an aggregate detectability for all checklists at a given location using 1- sum(1-$p_i$) where $p_i$ is the estimated detectability of wood thrush on a checklist. We selected all locations with an average detectability across all checklists in 2017 of at least 0.9. We estimated occupancy at these sites and compared this with whether wood thrush had been detected on any checklist at the site. The advantage of this dataset is that it allows us to test the occupancy submodel, without being confounded with the detectability submodel, however the disadvantage is that the validation is biased towards sites with several visits.

*Validation metrics*

We did not have a single criterion (e.g., accurate prediction of non-zero values, accurate prediction of rank order of observed values) with which to judge the predictive performance of models. Instead, we want to assess the extent to which any and all types of weaknesses exist. Given this objective, we have used multiple performance metrics, each of which emphasizes a different facet of model accuracy. For each model we selected a threshold for predicted presence that maximised Kappa and we used this for the metrics that require a threshold. We used a set of performance metrics available in the R packages *PresenceAbsence* (Freeman & Moisen, 2008) and *verification* (NCAR Research Applications Laboratory, 2015).

**References**

Dormann, C. F. (2020). Calibration of probability predictions from machine-learning and statistical models. *Global Ecology and Biogeography: A Journal of Macroecology*, 574392.

Fiske, I., & Chandler, R. (2011). unmarked: An R package for fitting hierarchical models of wildlife occurrence and abundance. *Journal of Statistical Software*, *43*(10), 1–23.

Freeman, E. A., & Moisen, G. (2008). PresenceAbsence: An R package for presence-absence model analysis. *Journal of Statistical Software*, *23*(11), 1–31.

James, G., Witten, D., Hastie, T., & Tibshirani, R. (2013). *An Introduction to Statistical Learning: with Applications in R*.

NCAR Research Applications Laboratory. (2015). *verification: Weather Forecast Verification Utilities*. Retrieved from https://CRAN.R-project.org/package=verification

Phillips, S. (2016). *Maxnet: Fitting "maxent"species distribution models with "glmnet."*

Pya, N. (2013). *scam: Shape constrained additive models*.

Sauer, J. R., & Link, W. A. (2011). Analysis of the North American breeding bird survey using hierarchical models. *The Auk*, *128*(1), 87–98.

Sauer, J. R., Pardieck, K. L., Ziolkowski, D. J., Smith, A. C., Hudson, M.-A. R., Rodriguez, V., … Link, W. A. (2017). The first 50 years of the North American Breeding Bird Survey. *The Condor*, *119*(3), 576–593.

Strimas-Mackey, M., Miller, E., & Hochachka, W. (2017). auk: eBird Data Extraction and Processing with AWK. *R Package Version*.

Wood, S. (2006). *Generalized Additive Models: An Introduction with R*. CRC Press.

Wright, M., & Ziegler, A. (2017). ranger: A Fast Implementation of Random Forests for High Dimensional Data in C++ and R. *Journal of Statistical Software, Articles*, *77*(1), 1–17.
