## Appendix A3 for "Analytical guidelines to increase the value of citizen science data: using eBird data to estimate species occurrence"

### SUPPORTING INFORMATION APPENDIX A3

#### General information for fitting species distribution models with eBird data

In this Appendix we describe in detail some of the more complex aspects of fitting the species distribution models with eBird data: accounting for variation in detectability among checklists, mitigating impacts of class imbalance, model validation, and defining the repeated visits for occupancy models. This is more general information about these models, rather than the specifics in the analyses in this paper. Further information and code to fit species distribution models in eBird data is available in the online bookdown document: “Best Practices for Using eBird Data” (Strimas-Mackey et al. 2020).

*Detectability*

With citizen science data it can be particularly important to consider detectability. In this context ‘detectability’ describes the probability that an individual or species that *occupies or uses* a given area will be detected, identified, and recorded by the participant. The detectability of birds by citizen scientists varies by ecological factors such as season, habitat, and species, in addition to observation factors such as time of day (Marques, Thomas, Fancy, & Buckland, 2007; Bas, Devictor, Moussus, & Jiguet, 2008; Lehikoinen, 2013; Kelling et al., 2015). There are two key aspects of detectability to consider in species distribution models; absolute detectability and variation in detectability. Absolute detectability can be estimated with occupancy modelling and under certain assumptions semi-structured citizen science data can be used to fit occupancy models (Kéry, Gardner, & Monnerat, 2010; Johnston, Fink, Hochachka, & Kelling, 2018). To account for variation in detectability, data can be filtered and covariates that describe the variation can be included in models. Projects that collect data describing the observation process will be able to account for a larger proportion of the heterogeneity in detectability (Kelling et al., 2018). The appropriate modelling framework depends on the goals of the analysis and the available data (Guillera-Arroita et al., 2015), but a greater variety of models are feasible when only estimating and accounting for variation in detectability.

*Class imbalance*

Class imbalance in citizen science data can be substantial, with a high proportion of checklists not reporting a species. This is particularly marked for species that are difficult to detect. Thus, looking for the degree of class imbalance should be standard practice when preparing to fit models to eBird data. The first step in this process is to determine the ratio of detection checklists to non-detection checklists for your species of interest.

One must consider class imbalance because not doing so may result in poorly performing models. As classes move away from equal balance, many classification methods become less accurate e.g. tree based methods, neural nets, Bayesian belief networks, support vector machines, k-nearest neighbor; (He & Garcia, 2009; Sun, Wong, & Kamel, 2009). Even simple models such as logistic regression lose predictive ability as the classes move away from balance (Fithian & Hastie, 2014). Another reason class imbalance should be addressed is that inference gleaned from models trained on imbalanced data may not be informative (Longadge & Dongre, 2013). For example, a distribution model for a species appearing on 1.5% of checklists could choose the default strategy of predicting that the species is not present anywhere across the spatial extent to which it predicts. This model would correctly predict whether the species was present or not roughly 98.5% of the time; very high accuracy. However, this model would not be very useful for trying to understand the habitat associations or distribution of the species.

There are many methods for dealing with unbalanced data. When using a random forest (as we do in the 'encounter probability' model), it is possible to use balanced random forests, wherein the model draws its bootstrap samples from each class separately, drawing first from the minority class, then drawing an equal number from the majority class (Chen, Tsai, Young, & Kodell, 2005; Agusta & Adiwijaya, 2018). Another method is weighted random forest, where the model assigns a harsher penalty to misclassification of the minority class than it does to the majority class (Chen et al., 2005; Agusta & Adiwijaya, 2018).

It is also possible to address class imbalance via a sampling routine before the modelling. One may use oversampling (by creating duplicated examples of the minority class), or undersampling (by randomly removing observations from the majority class) to create a more balanced dataset. Both techniques are often used together. One specific oversampling technique, synthetic minority oversampling technique (SMOTE) creates synthetic examples from the minority class. This creates synthetic examples that occupy the parameter space between randomly chosen observations and their nearest neighbors so that the added observations are not direct copies (Chawla, Bowyer, Hall, & Kegelmeyer, 2002).

When using eBird data it can be useful to jointly consider the spatial bias and the class imbalance. Spatial sampling can be combined with undersampling, and this method was shown to be very useful for eBird data (Robinson, Ruiz-Gutierrez, & Fink, 2018). This method targets a particular species and splits the data into two subsets; 1) checklists with species detections; and 2) checklists with non-detections for a given species. The non-detection dataset is then spatially filtered to address the spatial bias (see Supporting Information A2), and the filtered dataset is combined with the detection dataset. The result is a spatially undersampled data set for which the degree of class imbalance is mitigated. This method works well for species with a very low proportion of detections, as the spatial bias maintained by not spatially filtering the detection observations is minimal. It is also possible to spatially filter both datasets before recombining the filtered data sets (as we did to create the balanced validation set in this paper).

*Model Validation*

In order to assess the impacts of different practices on the accuracy of encounter-rate and occupancy models we used two semi-independent validation datasets. It is important to select validation data that is aligned with the objectives of the modelling exercise (Valavi, Elith, Lahoz-Monfort, & Guillera-Arroita, 2018). For example, is it more important to predict where species is present, or where species is absent? Is it important to get the ranking between sites correct, or the absolute values of occupancy? At what spatial and temporal scale do the models need to be relevant? Once the main objectives are defined, an appropriate validation dataset and metrics can be selected.

For our assessments we wanted to evaluate models based on their ability to produce accurate predictions across the entire spatial extent of our study region. For this purpose, it is appropriate to create a spatially balanced set of test data that evenly represents the study region. We also wanted to assess the ability of the models to discriminate between places of species presence and absence, so we created a balanced validation dataset with equal detections and non-detections. With the encounter rate models, the best way to do this is by assessing the ability of the models to discriminate between detections and non-detections in the validation dataset. See Appendix A2 for further details of the validation datasets used in this paper.

*Estimating species occupancy*

The initial stages of preparing data for occupancy modelling were the same as those described for modelling of encounter rates in the main paper. The data from each checklist were converted into binary detection/non-detection data during the process of creating zero-filled data, using the R package *auk*.

The next step in formatting data was to group our data into series of repeated count, with each series representing the data from a single 'site' that was repeatedly visited. We defined each site as being a combination of geographic point and observer (locality_id and observer_id; see Table A1). One other option would have been to allow multiple observers' data from the same geographic point to be combined into a single time set; however, in most cases all data from a locality still would have been from the same observer, so we opted for the most conservative approach of defining sites of combinations of geographic point and observer. The inclusion of the checklist_calibration_index (Table A1) as a site-level predictor in our models enabled much of the observer-linked variation in detection rates to be statistically removed, so that effectively the remaining site-level variation represented variation in the environments among geographic points. Another alternative definition for 'site' would have been to create a spatial grid across the study region, and treat all checklists within a grid cell as replicate counts. However, this space-for-time substitution is not universally appropriate (Kendall & White, 2009), and thus we chose not to have to deal with this added interpretational complexity.

The definition of which checklists can be used to form series of repeated visits to a site requires an explicit definition of a period of closure during which occurrence at a site is assumed not to change (i.e. the species is either always present or always absent). Prior knowledge indicates that the month of June represents a period of time after spring migration, and during the period of time during which Wood Thrushes are nesting. Thus, we defined the period of closure to be the entire month of June.

A final aspect of defining the time period of closure is the decision of how to deal with the existence of multiple years' observations within our data. We treated each calendar year as a separate temporal entity, because we could not assume that sites occupied in one calendar year would also be occupied in other years. Within each site and calendar year we only retained data if a site-year combination had data from at least two repeated visits. Where there were more than 10 visits, we randomly selected 10 of these. We also allowed separate years' data from the same site to appear in the data set (a process sometimes referred to as "stacking"). The R package *auk* was used to convert the input data from their original format into a form in which each series of observations from a site and year are treated as a separate row of data, using the *auk* function *format_unmarked_occu*.

Next we spatially subsampled the data that had been formatted for occupancy modelling as described in the previous paragraph, in order to create a final data set in which the density of data was distributed relatively evenly across the study region. We retained only a single randomly-chosen 'site' (i.e. a set of observations from a single location, observer and calendar year) within each 5 km hexagonal grid cell as described in Appendix A2.

**References**

Agusta, Z. P., & Adiwijaya, A. (2018). Modified balanced random forest for improving imbalanced data prediction. *International Journal of Advances in Intelligent Informatics*. doi:10.26555/ijain.v5i1.255

Bas, Y., Devictor, V., Moussus, J.-P., & Jiguet, F. (2008). Accounting for weather and time-of-day parameters when analysing count data from monitoring programs. *Biodiversity and Conservation*, *17*(14), 3403–3416.

Chawla, N. V., Bowyer, K. W., Hall, L. O., & Kegelmeyer, W. P. (2002). SMOTE: Synthetic Minority Over-sampling Technique. *Journal of Artificial Intelligence Research*, *16*, 321–357.

Chen, J. J., Tsai, C. A., Young, J. F., & Kodell, R. L. (2005). Classification ensembles for unbalanced class sizes in predictive toxicology. *SAR and QSAR in Environmental Research*, *16*(6), 517–529.

Fithian, W., & Hastie, T. (2014). Local case-control sampling: Efficient subsampling in imbalanced data sets. *The Annals of Statistics*. doi:10.1214/14-aos1220

Guillera-Arroita, G., Lahoz-Monfort, J. J., Elith, J., Gordon, A., Kuiala, H., Lentini, P. E., … Wintle, B. A. (2015). Is my species distribution model fit for purpose? Matching data and models to applications. *Global Ecology and Biogeography: A Journal of Macroecology*, *24*(3), 276–292.

He, H., & Garcia, E. A. (2009). Learning from Imbalanced Data. *IEEE Transactions on Knowledge and Data Engineering*, *21*(9), 1263–1284.

Johnston, A., Fink, D., Hochachka, W. M., & Kelling, S. (2018). Estimates of observer expertise improve species distributions from citizen science data. *Methods in Ecology and Evolution / British Ecological Society*, *9*, 88–97.

Kelling, S., Johnston, A., Fink, D., Ruiz-Gutierrez, V., Bonney, R., Bonn, A., … Guralnick, R. (2018). Finding the signal in the noise of Citizen Science Observations. *bioRxiv*. Retrieved from https://www.biorxiv.org/content/early/2018/05/18/326314.abstract

Kelling, S., Johnston, A., Hochachka, W. M., Iliff, M., Fink, D., Gerbracht, J., … Yu, J. (2015). Can Observation Skills of Citizen Scientists Be Estimated Using Species Accumulation Curves? *PloS One*, *10*(10), e0139600.

Kendall, W. L., & White, G. C. (2009). A cautionary note on substituting spatial subunits for repeated temporal sampling in studies of site occupancy. *The Journal of Applied Ecology*, *139*, 657.

Kéry, M., Gardner, B., & Monnerat, C. (2010). Predicting species distributions from checklist data using site-occupancy models. *Journal of Biogeography*, *37*(10), 1851–1862.

Lehikoinen, A. (2013). Climate change, phenology and species detectability in a monitoring scheme. *Population Ecology*, *55*(2), 315–323.

Longadge, R., & Dongre, S. (2013, May 8). *Class Imbalance Problem in Data Mining Review. arXiv [cs.LG]*. Retrieved from http://arxiv.org/abs/1305.1707

Marques, T. A., Thomas, L., Fancy, S. G., & Buckland, S. T. (2007). Improving estimates of bird density using multiple-covariate distance sampling. *The Auk*, *124*(4), 1229–1243.

Robinson, O. J., Ruiz-Gutierrez, V., & Fink, D. (2018). Correcting for bias in distribution modelling for rare species using citizen science data. *Diversity & Distributions*, *24*(4), 460–472.

Strimas-Mackey, M., W.M. Hochachka, V. Ruiz-Gutierrez, O.J. Robinson, E.T. Miller, T. Auer, S. Kelling, D. Fink, A. Johnston. (2020). Best Practices for Using eBird Data. Version 1.0. https://cornelllabofornithology.github.io/ebird-best-practices/. Cornell Lab of Ornithology, Ithaca, New York. https://doi.org/10.5281/zenodo.3620739

Sun, Y., Wong, A. K. C., & Kamel, M. S. (2009). CLASSIFICATION OF IMBALANCED DATA: A REVIEW. *International Journal of Pattern Recognition and Artificial Intelligence*, *23*(04), 687–719.

Valavi, R., Elith, J., Lahoz-Monfort, J. J., & Guillera-Arroita, G. (2018). block CV: An r package for generating spatially or environmentally separated folds for k -fold cross-validation of species distribution models. *Methods in Ecology and Evolution / British Ecological Society*, *67*, 617.
