## Supplementary Figures for "Analytical guidelines to increase the value of citizen science data: using eBird data to estimate species occurrence"

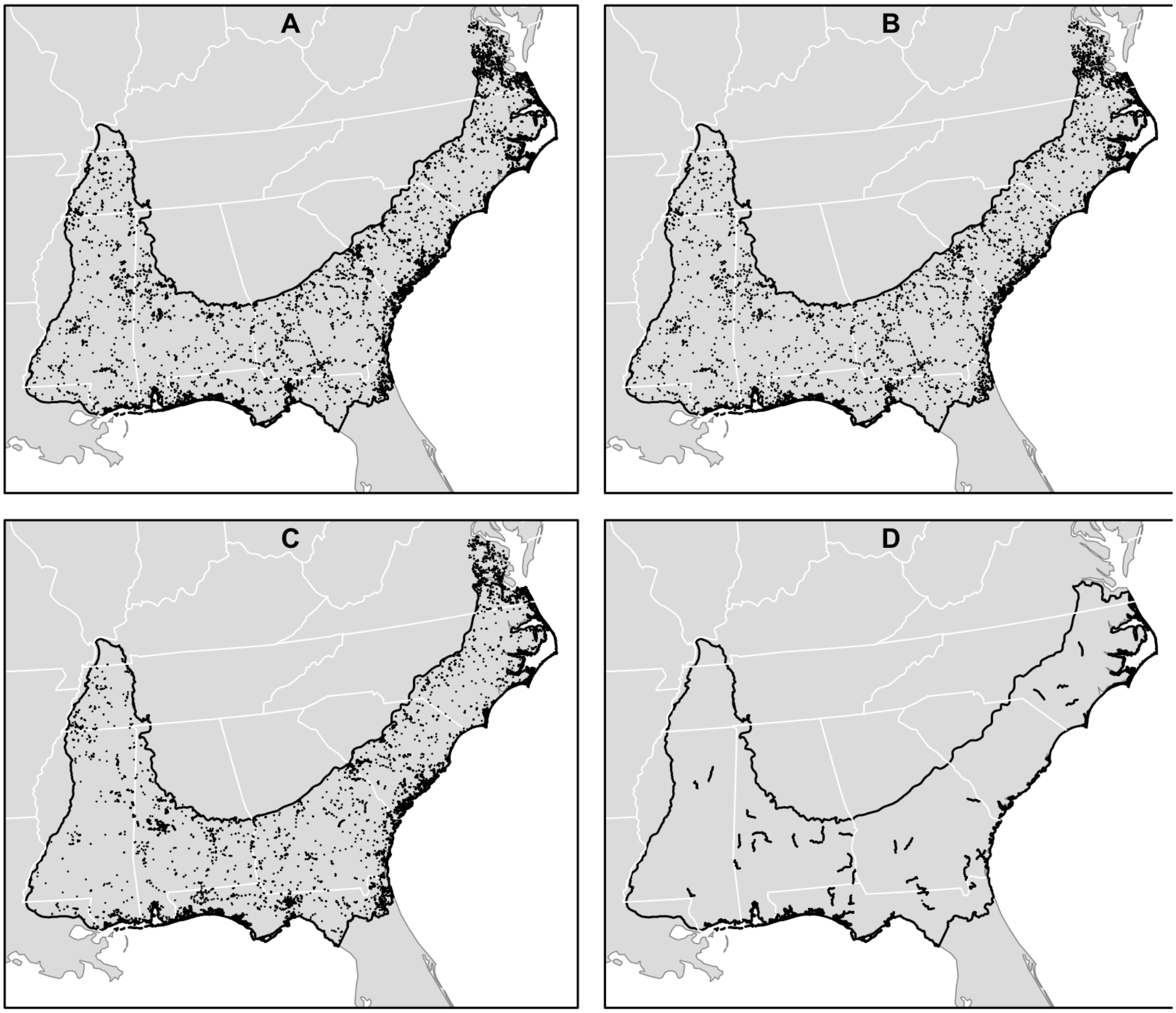


**Figure S1.** Location of checklists for A all training data 2018; B example of training data selecting only complete checklists and then spatially subsampled (data for Model 5); C test data 2017; D test data eBird-entered BBS data 2018.

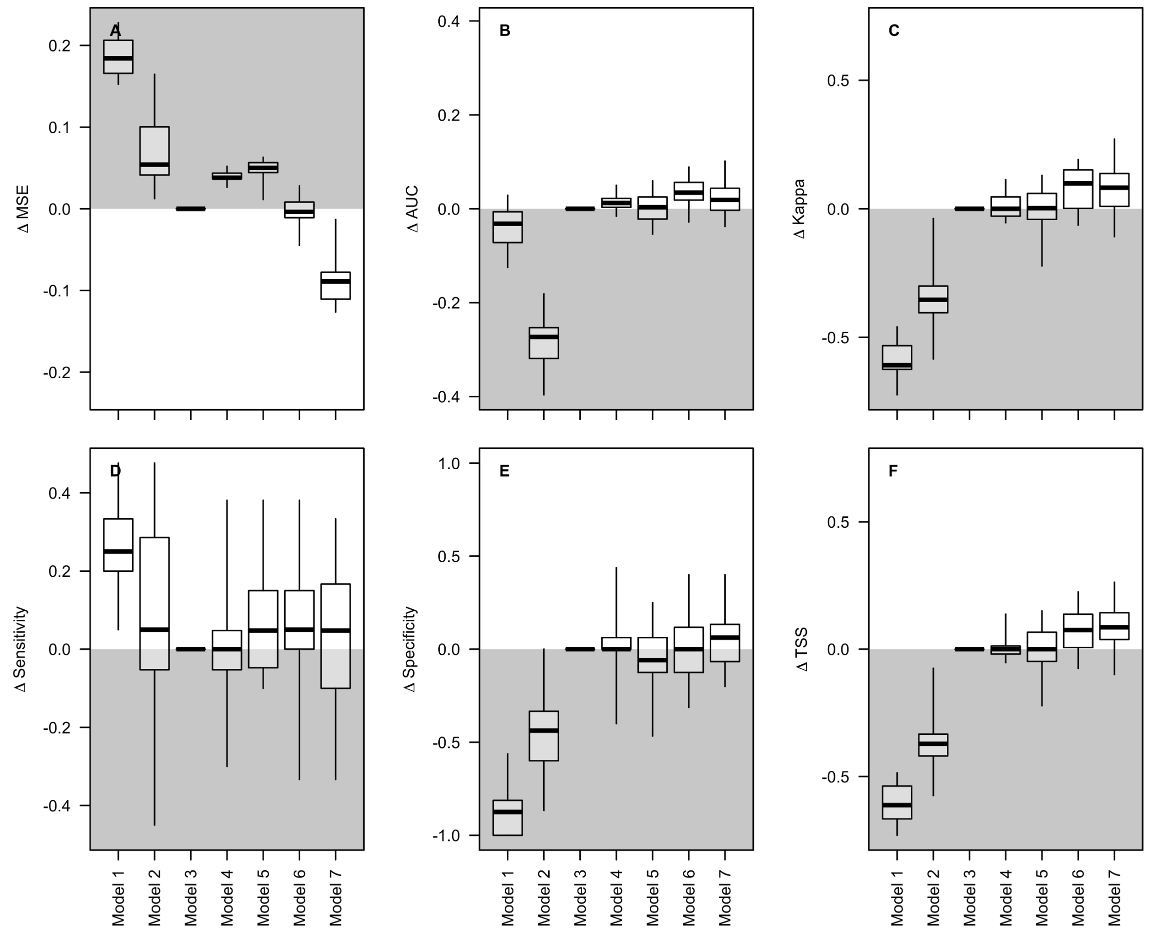


**Figure S2.** Differences in predictive performance metrics for the encounter rate models 1-7 against BBS data within eBird from 2018, aggregated to BBS routes. Metrics are compared to the performance from model 3, with the y-axis values showing the differences relative to model 3. The white halves of the plots indicate where model performance is *better* than model 3. The grey halves of the plots indicate where model performance is *worse* than model 3. Model 1 is the Maxent model which uses only presences and produces background psuedo-absences. Model 7 is the random forest encounter rate model using complete checklists, spatial subsampling, effort variable filters, and effort variables as covariates. The validation metrics are calculated for 25 different model runs. Further details of models 1-7 are in the text and Table 1. Boxes show the median, the interquartile range, and the extremes of the distributions.

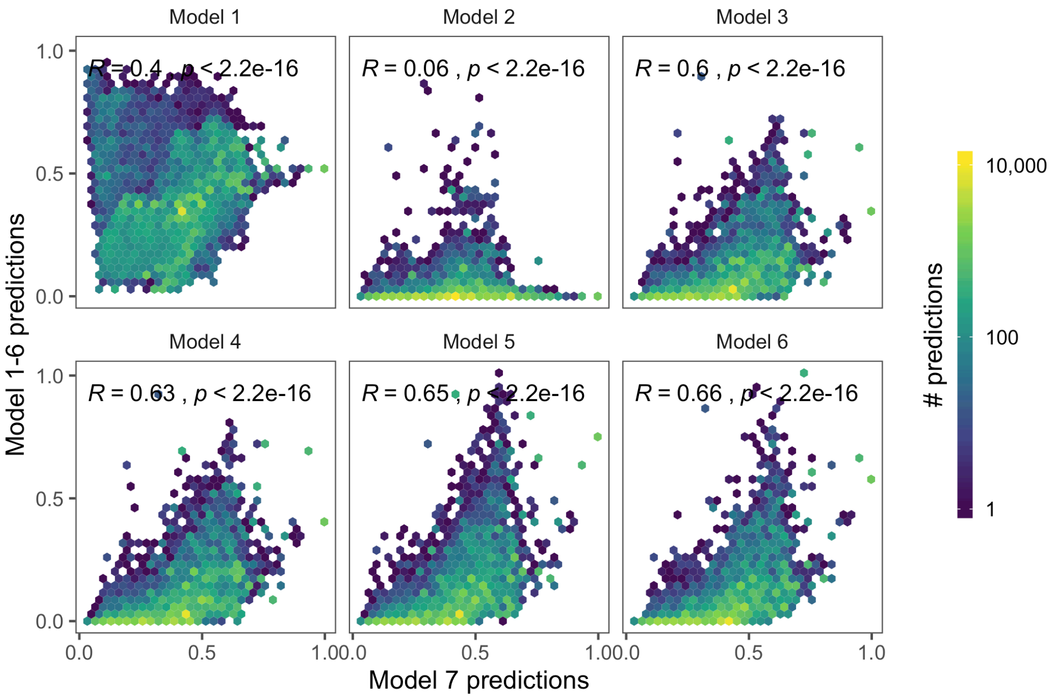


**Figure S3.** Density scatterplots comparing estimates of encounter rate across the BCR27 region for models 1 - 6 against model 7. Brighter colours indicate a higher density of points in that region of the scatterplot. The high densities along the bottom of the figures represent many locations that are predicted to have low encounter rate with models 1-6 and higher encounter rate with model 7. R values show the Pearson correlation coefficient and the associated p-value.

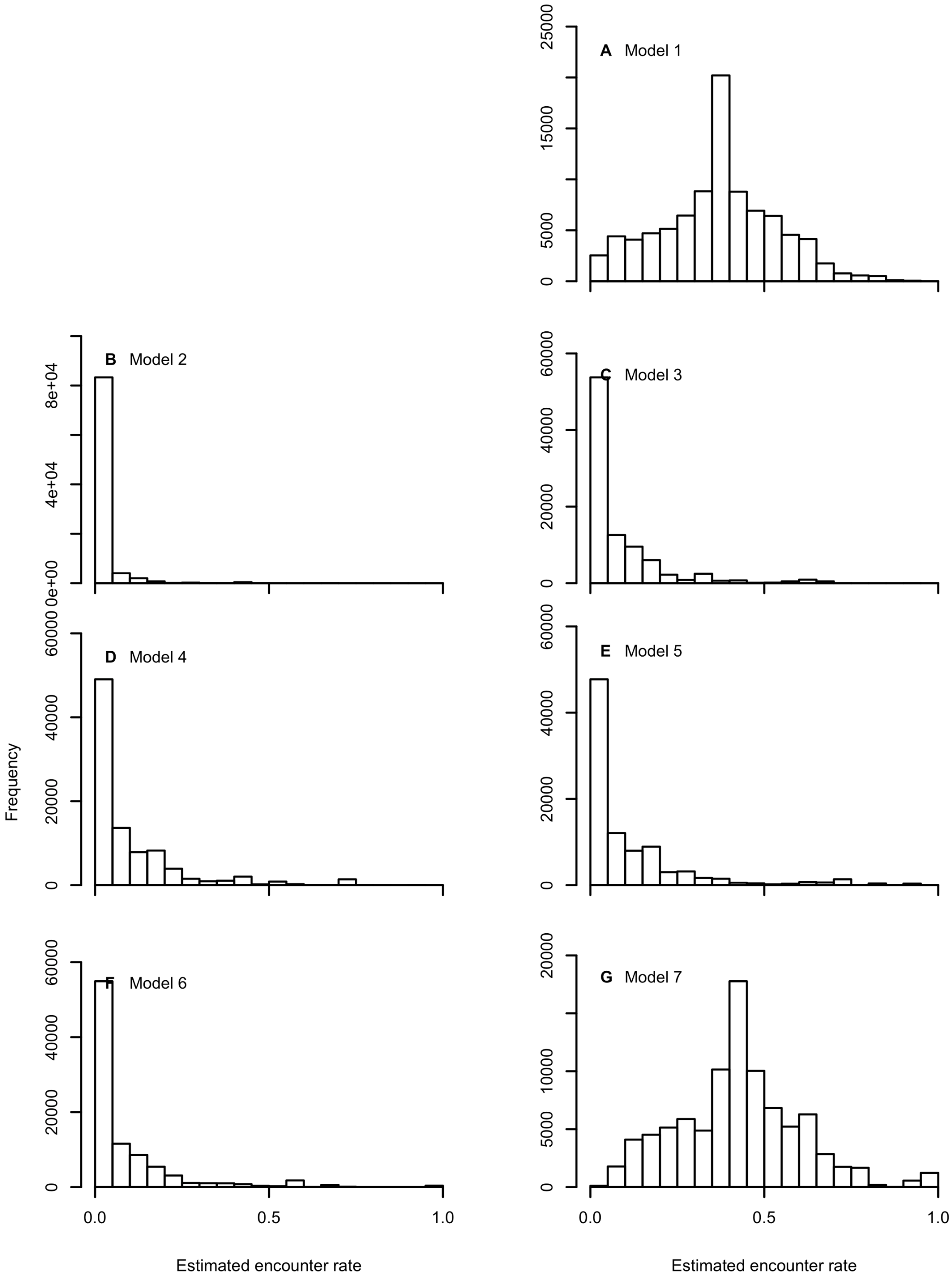


**Figure S4.** Estimated encounter rates across BCR27 for models 1-7. See Table 1, Figure 1, Appendix S2 and the main text for full details of each model.

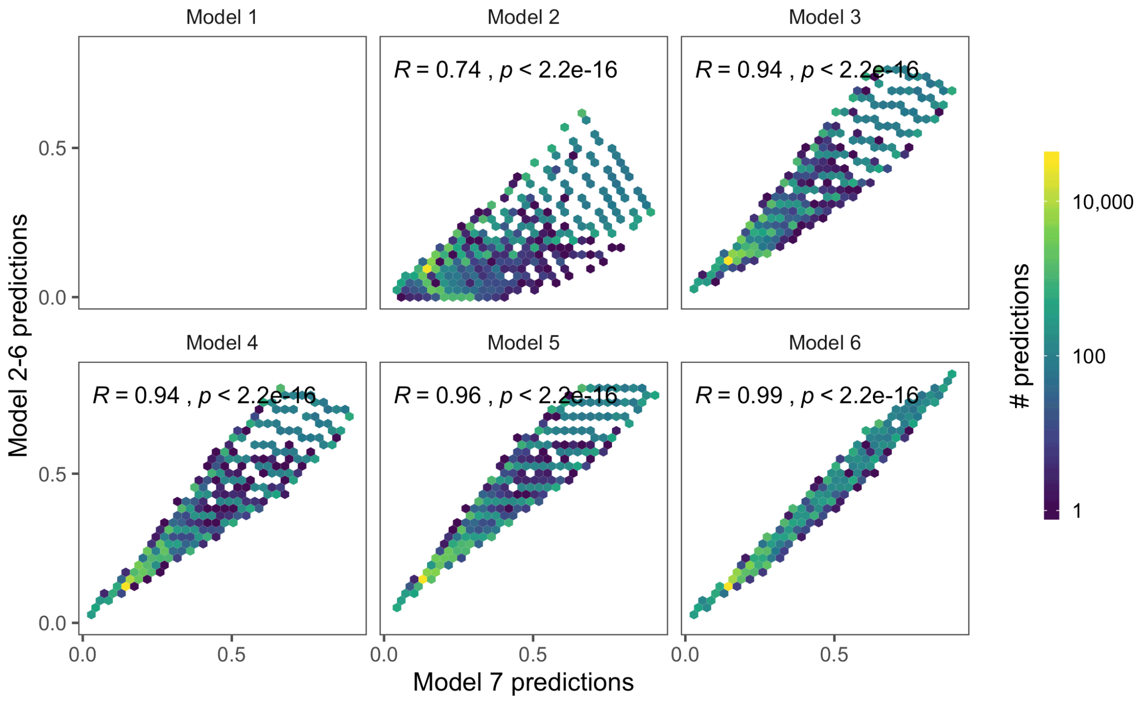


**Figure S5**. Density scatterplots comparing estimates of occupancy rate across the BCR27 region for models 2 - 6 against model 7. Brighter colours indicate a higher density of points in that region of the scatterplot. R values show the Pearson correlation coefficient and the associated p-value.

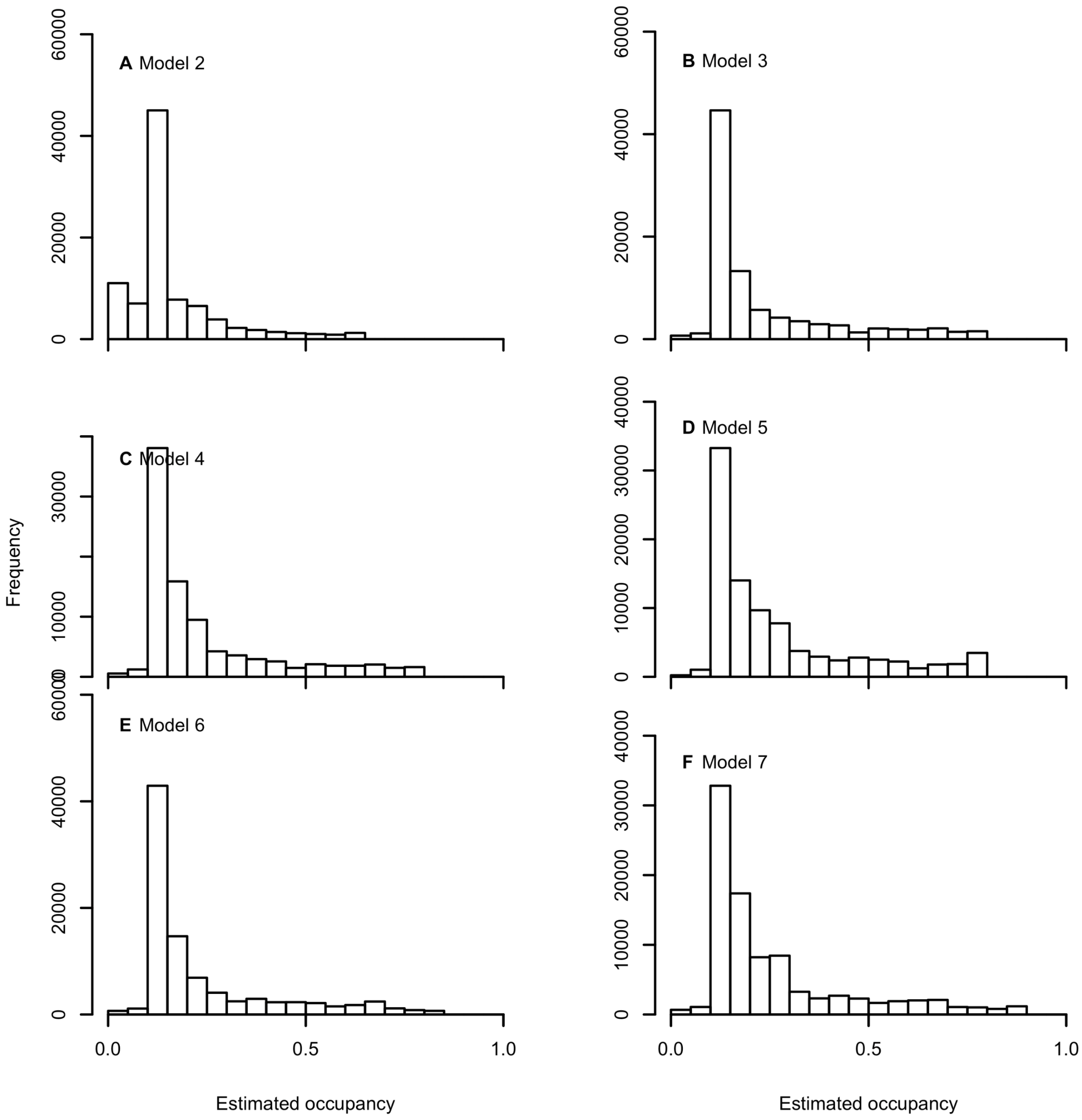


**Figure S6**. Estimated occupancy rates across BCR27 for models 2-7. See Table 1, Figure 1, Appendix S2 and the main text for full details of each model.

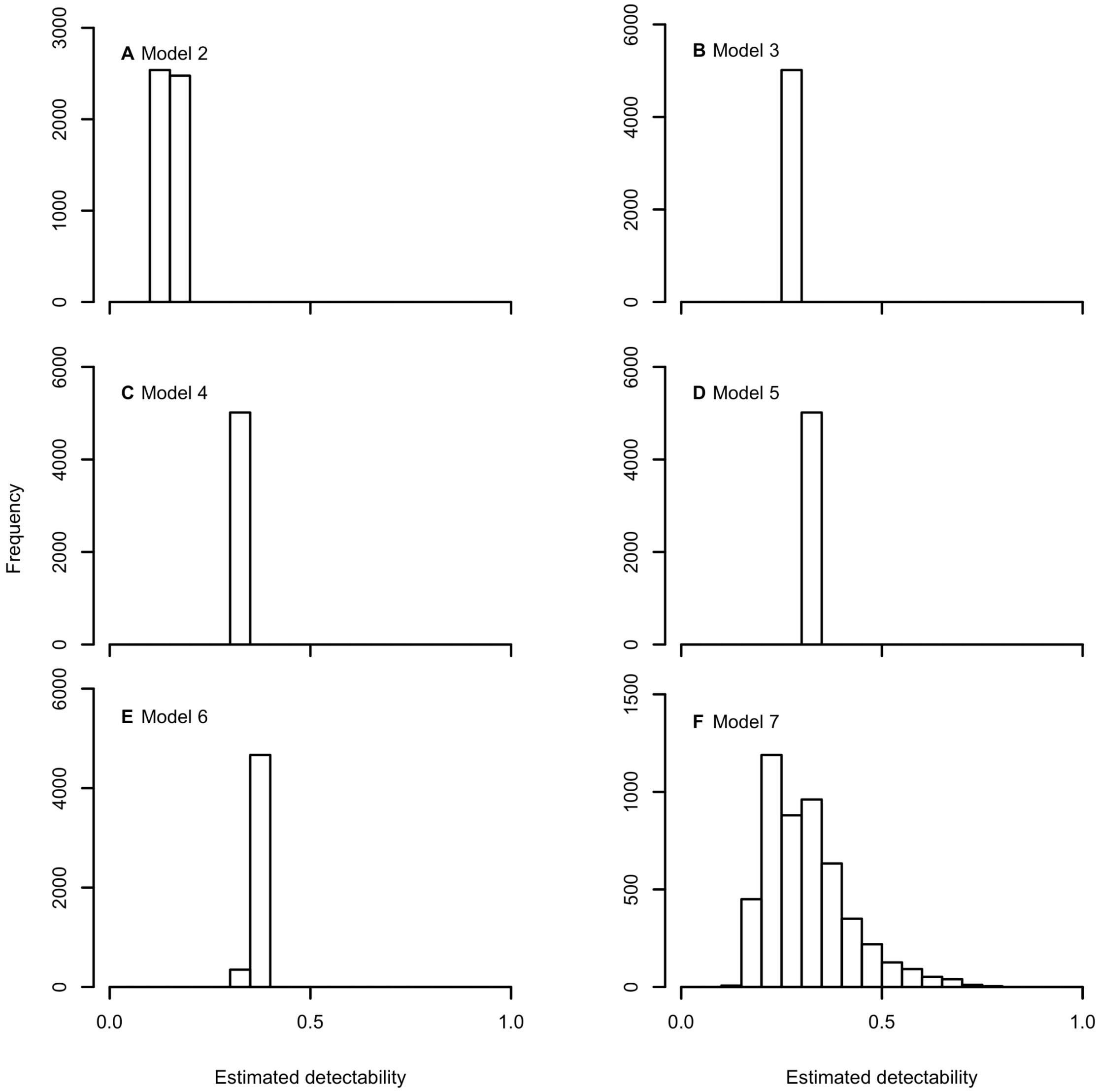


**Figure S7**. Estimated detectability rates in the training data used for model 7, but predicted from models 2-7. See Table 1, Figure 1, Appendix S2 and the main text for full details of each model.

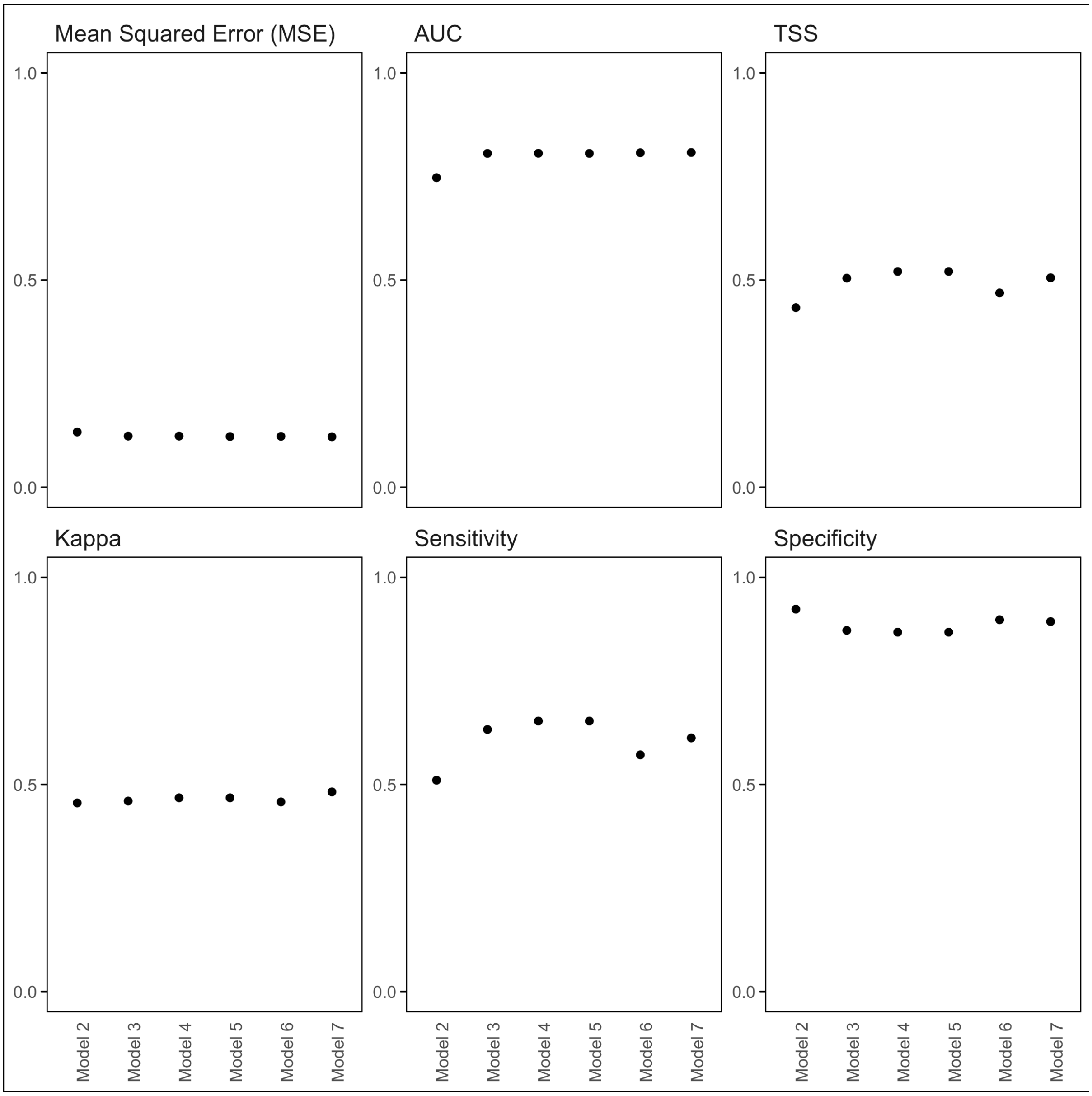


**Figure S8**. Performance metrics for occupancy models 2-7, created with 2018 data and validated with well-surveyed sites in 2017.

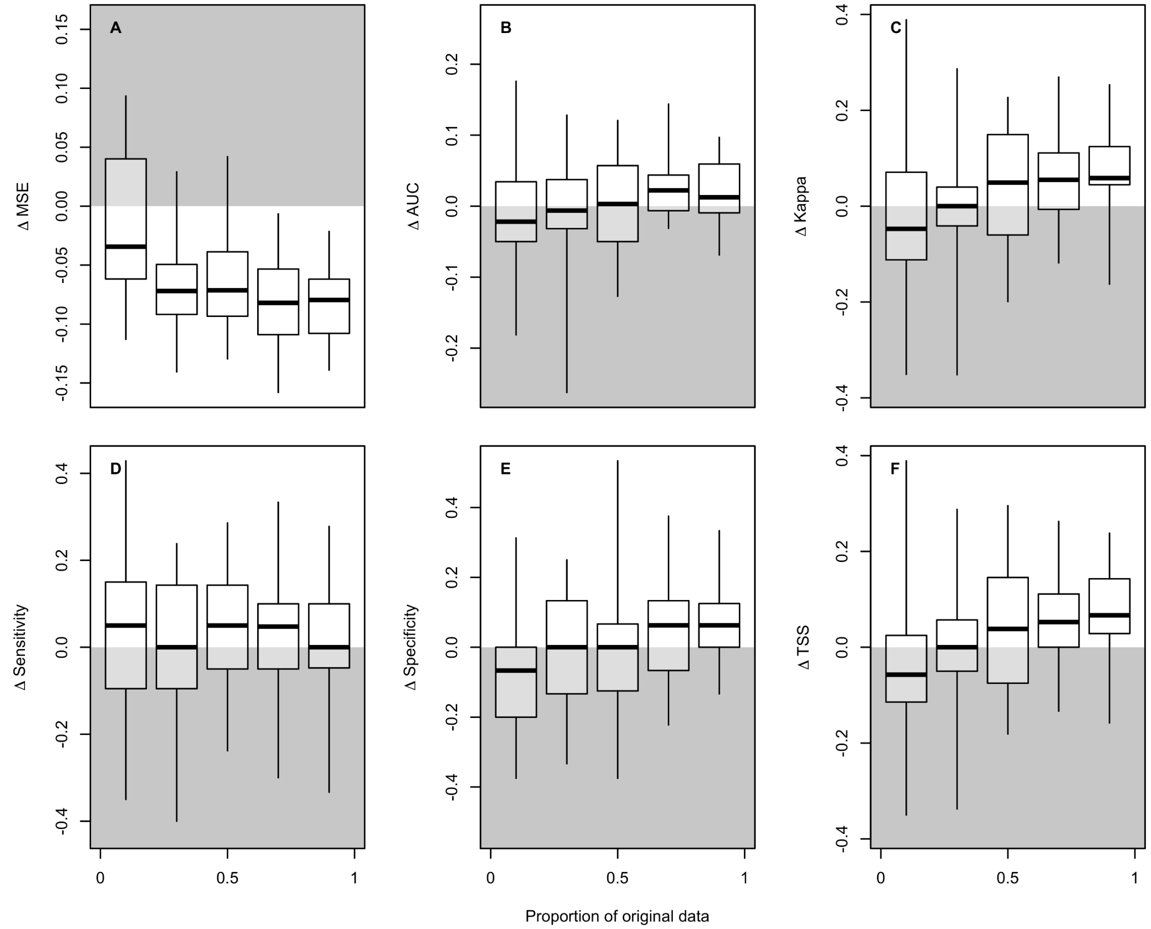


**Figure S9.** Effect of sample size on differences in predictive performance metrics for the encounter rate models 3 and 7. Differences were computed between the metrics as (Model 7 - Model 3). The white half of the plots indicating where Model 7 performance is *better* than Model 3. The grey half of the plots indicating where Model 7 performance is *worse* than Model 3. The test dataset was BBS data within eBird from 2018, aggregated to routes. The datasets were random subsampled to 0.75 of the original checklists. Then they were further reduced to a proportion of this dataset: 0.1, 0.3, 0.5, 0.7, 0.9. This process was repeated 25 times to produce 25 paired comparisons of model performance for each dataset size. Each paired comparison between Model 3 and Model 7 used the same randomly subsampled test and train datasets. See Table 1 for further details of Model 3 and Model 7. Panels show the following performance metrics: A Mean Squared Error (MSE); B Area Under the Curve (AUC); C Kappa; D sensitivity; E specificity; and F True Skill Statistic (TSS). Boxes show the median, the interquartile range, and the extremes of the distributions.
